# IGREX for quantifying the impact of genetically regulated expression on phenotypes

**DOI:** 10.1101/546580

**Authors:** Mingxuan Cai, Lin S. Chen, Jin Liu, Can Yang

## Abstract

By leveraging existing GWAS and eQTL resources, transcriptome-wide association studies (TWAS) have achieved many successes in identifying trait-associations of genetically-regulated expression (GREX) levels. TWAS analysis relies on the shared GREX variation across GWAS and the reference eQTL data, which depends on the cellular conditions of the eQTL data. Considering the increasing availability of eQTL data from different conditions and the often unknown trait-relevant cell/tissue-types, we propose a method and tool, IGREX, for precisely quantifying the proportion of phenotypic variation attributed to the GREX component. IGREX takes as input a reference eQTL panel and individual-level or summary-level GWAS data. Using eQTL data of 48 tissue types from the GTEx project as a reference panel, we evaluated the tissue-specific IGREX impact on a wide spectrum of phenotypes. We observed strong GREX effects on immune-related protein biomarkers. By incorporating trans-eQTLs and analyzing genetically-regulated alternative splicing events, we evaluated new potential directions for TWAS analysis.

## Introduction

Genome-wide association studies (GWAS) have successfully identified tens of thousands of unique associations between single-nucleotide polymorphisms (SNPs) and a wide range of complex traits/diseases (http://www.ebi.ac.uk/gwas/). More than 90% of identified risk variants are located in non-coding regions [1], making it challenging to understand their functional mechanisms. Increasing evidence [2, 3, 4, 5, 6, 7, 8, 9] has suggested that many of those risk variants may affect traits/diseases via the modulation of their cis gene expression levels. For example, a study of 18 complex traits revealed an enrichment for expression quantitative trait loci (eQTLs) in 11% of 729 tissue-trait pairs [10]. There is great interest in precisely characterizing the specific role of genetically regulated gene expression (GREX) in human traits and diseases.

It is well known that the effects of genetic variation on gene expressions depend on cellular contexts [11]. The rapidly increasing availability of eQTL data from different tissue types, cell types, populations and other conditions provides an unprecedented opportunity to study and evaluate GREX effects in a variety of conditions. For example, the V7 release of the Genotype-Tissue Expression (GTEx) project (https://gtexportal.org/home/) has collected gene expression samples from 53 non-diseased tissues across 714 individuals [11]. Multiple blood eQTL resources comprising thousands of individuals are made publicly available [12, 13]; and other ongoing projects such as Genetics of DNA Methylation Consortium (GoDMC) and eQTLGen consortium are collecting expression data with sample sizes larger than 10,000 [14, 15]. Those data serve as rich eQTL resources for a comprehensive evaluation of GREX effects.

The vast amount of publicly available eQTL and GWAS data resources enables an integrative framework, transcriptome-wide association studies (TWAS), for mapping gene-level trait associations and evaluating GREX effects on human traits and diseases. Using a reference eQTL panel (e.g., GTEx), gene-specific expression prediction models can be built based on cis-acting genetic factors. Then the gene expression levels of a GWAS cohort can be predicted based on individual genetic profiles, and the genetically-regulated and predicted expression levels are further associated with the phenotype of interest in the GWAS study to map gene-level trait-associations [16, 17, 18, 19, 20, 21, 22, 23, 24]. Existing methods have been proposed [8, 25], including PrediXcan [16], TWAS [17], FOCUS [19], S-PrediXcan [21], UTMOST [26] and CoMM [22]. Through applications to a wide variety of phenotypes, these methods have successfully identified specific gene-trait associations, whereas a comprehensive and precise evaluation of the impact of GREX variation on various traits and the trait-relevant cellular context is still needed [27].

TWAS-types of integrative analysis rely on a key assumption: there exists a steady-state GREX variation shared across reference eQTL data and GWAS data, and the steady-state GREX variation can further induce phenotypic variation. The multi-tissue eQTL data from the GTEx project is commonly used as the reference eQTL panel [16, 21, 26]. The GTEx project has collected data from post-mortem donors and has provided a source of largely non-diseased tissues for general purposes. The GTEx reference may or may not have considerable shared GREX variation with GWAS data of specific phenotypes in specific populations. Given the often unknown disease/trait-relevant tissue types and the increasing availability of eQTL data resources from different conditions, there is a need for new methods and tools that can be used to assess the proportion of the shared GREX variation in the phenotypic variation from a global perspective, and guide the selection of eQTL reference data and tissue-types for specific phenotypes and populations.

The heritability measure has been widely used to quantify the impact of genetic variation on phenotypic variation, and has served as a preliminary yet insightful assessment of the potential of genetic studies on various phenotypes [28, 29]. Analogous to the heritability measure, the estimation of proportion of GREX on phenotypic variation can also be used to evaluate the impact of the genetic regulatory effects on phenotypes mediated by expression levels, and inform trait-relevant tissue types or conditions in specific populations. To the best of our knowledge, there are two methods that have been proposed for this purpose [23, 24]. The RhoGE method [23] estimates the proportion of phenotypic variation explained by GREX based on linkage-disequilibrium (LD) score regression (LDSC) [30]. Since it ignores the uncertainty in predicting gene expression levels, the proportion of variance explained by GREX could be substantially under-estimated by RhoGE. The other method, known as the gene expression co-score regression (GECS) [24], requires the analyzed SNPs not being in LD to ensure unbiasedness, which greatly limits its applicability in real data analysis.

In this work, we propose a unified framework, named IGREX, for quantifying the impact of genetically regulated expression, while accounting for uncertainty in predicted gene expression levels in the presence of moderate to weak eQTL effects. IGREX requires only summary-level GWAS data as input, greatly enhancing the applicability of the method. We evaluated the performance of IGREX with comprehensive simulation studies, highlighting the importance of accounting for expression estimation uncertainty. Using 48 tissue types from the GTEx project as the reference panel, we applied IGREX to both individual-level and summary-level GWAS data sets, and evaluated the tissue-specific IGREX impact on a wide spectrum of cellular and organismal phenotypes. Our results provide new biological insights into the role of gene expression in the genetic architecture of complex traits. We also demonstrate the reproducibility of results. By incorporating trans-eQTLs and analyzing genetically-regulated alternative splicing events, we evaluated new potential directions for TWAS analysis.

## Results

### Method overview

IGREX is a two-stage method for quantifying the proportion of phenotypic variation that can be attributed to GREX variation. The method can be applied to both individual-level (IGREX-i) and summary-level (IGREX-s) GWAS data. It first evaluates the posterior distribution of GREX effects based on an eQTL reference panel and then estimates the proportion of variance explained by GREX using the ‘predicted’ gene expression in the GWAS data. Here, we briefly introduce the statistical model of IGREX-i and present additional technical details in the Methods Section.

Consider a reference eQTL data set 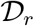 and an individual-level GWAS data set 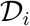. The eQTL data 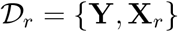 is comprised of an *n_r_* × *G* gene expression matrix, **Y**, and an *n_r_* × *M* genotype matrix, **X**_*r*_, where *G* is the number of genes, *M* is the number of SNPs and *n_r_* is the sample size. The GWAS data 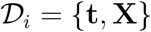 contains a phenotype vector **t** ∈ ℝ^*n*^ and a genotype matrix **X** ∈ ℝ^*n*×*M*^, where *n* is the sample size of the GWAS data. Let **y**_*g*_ and ***X***_*r,g*_ be the vector of expression levels of the *g*-th gene and the genotype matrix corresponding to its local (cis) SNPs from the reference panel, respectively. We first relate **y**_*g*_ to **X**_*r,g*_ with a linear model:

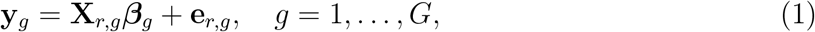

where 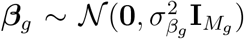 is the vector of genetic effects of *M_g_* cis SNPs on the expression levels of the *g*-th gene, and 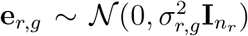 is the error term. Since we are interested in the steady-state component of gene expression levels regulated by genetic variants, *β_g_* is assumed to be the same for individuals in both datasets, 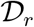 and 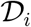. Consequently, the GREX component of individuals in the GWAS data can be evaluated by **X**_*g*_***β**_g_*. Meanwhile, we assume that the genetic effects on the phenotype of interest **t** can be decomposed into two parts, i.e. the effects mediated via GREX and the effects through alternative pathways not mediated by gene expression levels:

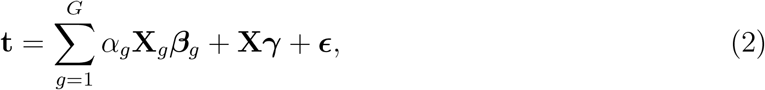

where 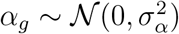 is the effect size of **X**_*g*_***β***_*g*_ on **t**, 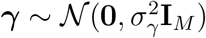 is the vector of alternative genetic effects, and 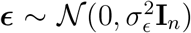 is the error term. In this model, 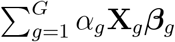 and **X_*γ*_** correspond to the overall impact of the GREX component and the alternative genetic effects on **t**, respectively. Thus, the impact of GREX on the phenotype can be quantified by the proportion of phenotypic variance explained by the GREX component: 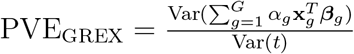.

To estimate this quantity, we propose a two-stage procedure: In the first stage, we estimate 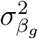 and 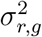 using an efficient algorithm and evaluate the posterior distribution 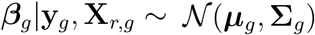 for all genes. In the second stage, by treating the posterior obtained in the first stage as the prior distribution of ***β**_g_* in model (2), we can obtain estimated values of 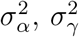 and 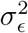 using either method of moments (MoM) or restricted maximum likelihood (REML). Following this procedure, the resulting estimate of PVE_GREX_ is obtained (with details in the Methods Section) by

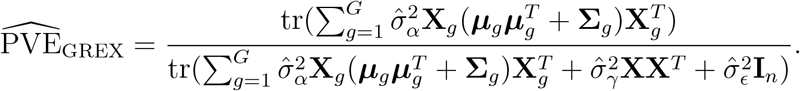

In the above estimation, the substitution of posterior ***β**_g_* |**y**_*g*_, **X**_*r,g*_ accounts for the posterior variance **Σ**_*g*_ and naturally results in the adjustment of estimation uncertainty associated with ***β***_*g*_. This is important because in the GWAS data, the gene expression levels are not directly measured, but rather are predicted or imputed based on genetic variants. It is known that the prediction accuracy and uncertainty vary substantially among genes. For most of the genes in the genome, the genetically regulated expression variation accounts for only a small to moderate proportion of total expression variation. Thus, the prediction may not be accurate and could be subject to high uncertainty. In contrast, our model accounts for the estimation uncertainty by **Σ**_*g*_ and can yield unbiased estimation for 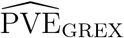. In addition, the standard error of 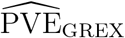 can be obtained based on the delta method (see Supplementary Note). The IGREX framework can also be used to test *H*_0_: PVE_GREX_ = 0 for the phenotype of interest in specific populations given an eQTL reference with a specific tissue type or cellular context.

In real applications, individual-level GWAS data may not be accessible. We have further developed IGREX-s which requires only summary-level GWAS data as input (See Methods). Based on MoM, IGREX-s can approximate IGREX-i while requiring only SNP-level *z*-scores from GWAS and a reference genotype matrix 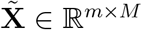 of a similar LD pattern to **X**, where *m* is the number of samples in the reference panel. Using simulations, we showed that with a few hundreds of samples in the eQTL reference data, the estimation of IGREX-s with summary statistics well approximates IGREX-i using individual level data. In practice, 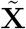 can be **X**_*r*_ or a subset of **X**. The estimate of PVE_GREX_ given by IGREX-s is

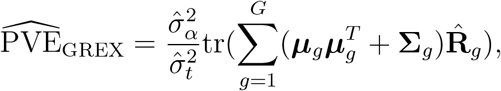

where 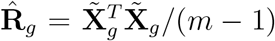 is the estimated LD matrix associated with the *g*-th gene and 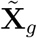 is the corresponding columns of 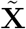. IGREX also allows for the adjustment of covariates including sex, age and genotype principal components (See details in Supplementary Note).

### Simulation studies

We conducted extensive simulation studies to evaluate the performance of IGREX. For all the simulated data, we fixed *n* = 4, 000, *G* = 200, *M* = 20,000 (i.e., 100 cis SNPs for each gene). The total phenotypic heritability was set as 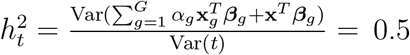, where PVE_GREX_ = 0.2 and the proportion explained by the alternative genetic effects, 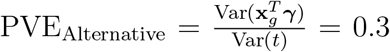 (results for other scenarios are shown in Supplementary Fig. 1-3). To simulate the genotype data, we first sampled the minor allele frequencies (MAF) from uniform distribution 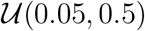 and data matrices from normal distribution 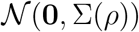, where Σ_*jj′*_ = *ρ*^|*j−j′*|^ characterizes the LD patterns between SNPs. Then, the genotype matrices **X**_*r*_ and **X** were obtained by categorizing the entries of generated data matrices into 0, 1, 2 according to MAF. Given the genotype matrices, ***β**_g_* and *α_g_*, the gene expression **y**_*g*_ and phenotype **t** were simulated following models (1) and (2). We will discuss the details for generating ***β**_g_* and *α_g_* later. To assess IGREX-s, we calculated the *z*-score of each SNP and randomly subsetted *m* = 500 samples from **X** for estimating LD matrix 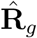 (results for other settings of *m* are shown in Supplementary Fig. 4).

We first evaluated the estimation performance of IGREX for different settings of eQTL reference data. Specifically, we varied *n_r_* at {800,1000, 2000}, 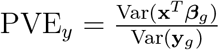 at {0.1, 0.2, 0.3}, where PVE_*y*_ quantifies the gene expression heritability explained by its local SNPs. To mimic the scenario in which the expression estimation uncertainty was incorrectly ignored, we obtained the posterior mean of ***β**_g_* in the first stage, and replaced the true effect size ***β**_g_* by its posterior mean ***μ**_g_* while specifying the posterior variance to be **Σ**_*g*_ = **0** in the second stage, and then conducted REML and MoM as before. We denoted these methods as REML_0_ and MoM_0_. The simulation results summarized in Fig.1a showed that both PVE_GREX_ and PVE_Alternative_ were accurately estimated using REML-based IGREX-i in all settings. The MoM-based IGREX-i slightly underestimated PVE_GREX_ when both sample size *n_r_* and PVE_*y*_ were very small, but steadily achieved similar performance as REML-based estimation when either *n_r_* or PVE_*y*_ increased. In all settings, IGREX-s well approximated MoM, producing nearly identical estimation results. In contrast, both REML_0_ and MoM_0_ did not account for estimation uncertainty in the expression prediction, and they showed poor estimation performance even when sample size was large and PVE_*y*_ value was high.

**Figure 1:**
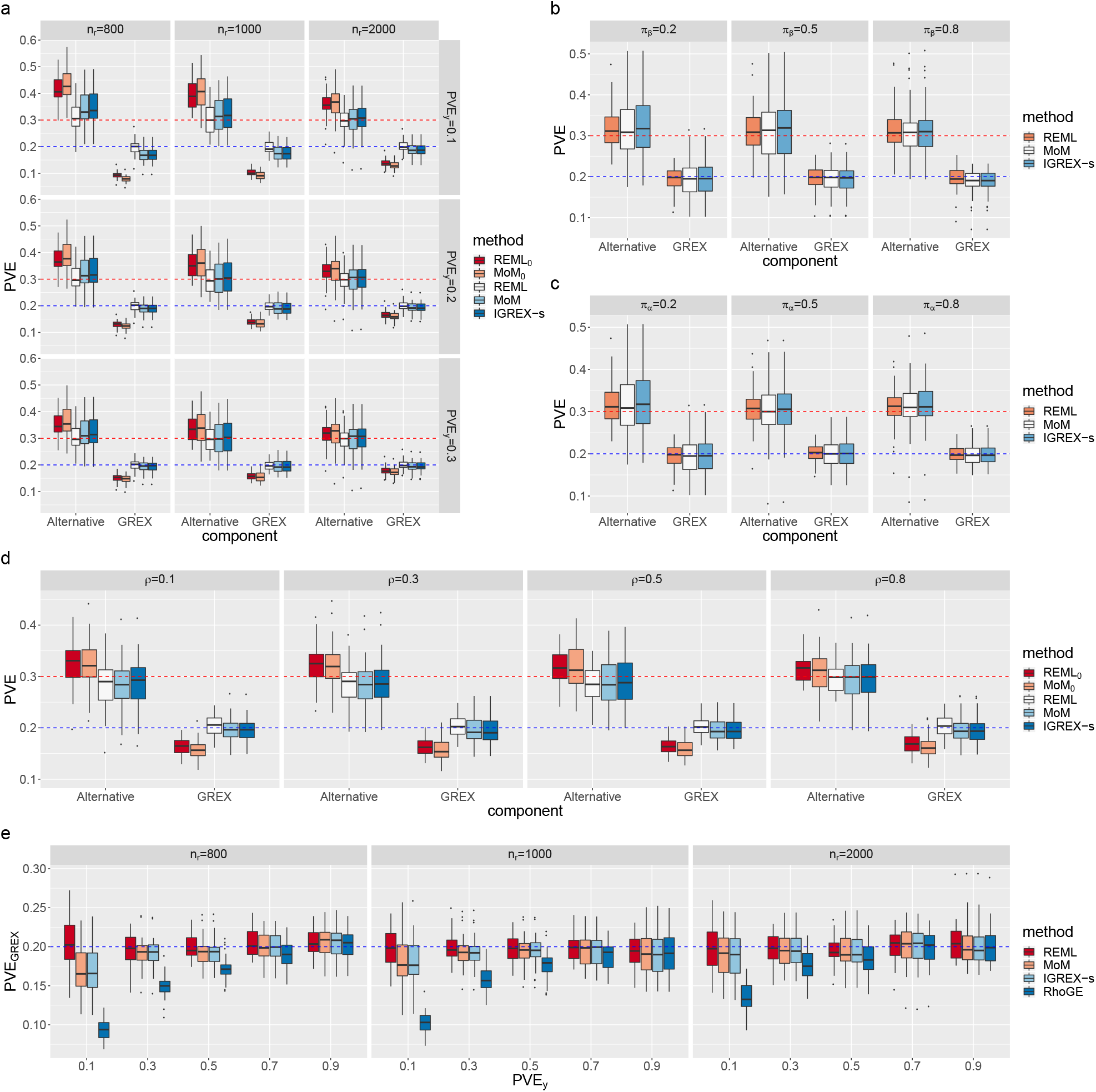
Simulation studies to compare estimation accuracies of IGREX with other methods. REML and MoM in the legend are abbreviations of the IGREX-i estimation methods. The blue and red dashed lines represent the true values of PVE_GREX_ and PVE_Alternative_, respectively. We averaged the results over 30 replications and generated box plots for evaluating the estimation performance of: **a** the three models of IGREX, REML_0_ and MoM_0_ when *n_r_* was varied at {800,1000, 2000} and PVE_*y*_ was varied at{0.1, 0.2, 0.3}; (**b**) the three models of IGREX when *π_α_* = 0.2 and *π_β_* was varied at {0.2, 0.5, 0.8}; (**c**) the three models of IGREX when *π_β_* = 0.2 and *π_α_* were varied at {0.2, 0.5, 0.8}; (**d**) the three models of IGREX, REML_0_ and MoM_0_ when *ρ* was varied at {0.1,0.3, 0.5,0.8}; (**e**) the three models of IGREX and RhoGE when *n_r_* was varied at {800,1000, 2000}.

Next we conducted simulations to evaluate the situation that the IGREX model was mis-specified. Here we considered the situation where genetic effects ***β**_g_* and ***α*** were sparse while we assumed dense effect sizes in the IGREX model. This was designed to mimic the real data situation that the architecture of eQTL signals is often sparse [31]. Let *π_α_* and *π_β_* be the sparsity of ***α*** and ***β**_g_*, i.e., *π_α_* = (# Nonzero entries in ***α***)/*G* and *π_β_* = (# Nonzero entries in ***β**_g_*)/*M_g_*, respectively. To evaluate the influence of different sparsity patterns on our method, we varied *π_α_* and *π_β_* at {0.2, 0.5, 0.8}. The nonzero entries in ***α*** and ***β***_*g*_ were simulated form a normal distribution. As shown in Fig. 1b-c, all three methods of IGREX produced accurate estimates in the presence of sparse genetic effects, implying the robustness of IGREX to model mis-specification. Moreover, the estimation performance was not influenced by the degree of sparsity. Next, we investigated the influence of LD patterns by letting *ρ* vary at {0.1, 0.3,0.5, 0.8}. From Fig.1d, we observed that IGREX produced accurate estimation in the presence of LD. In contrast, REML_0_ and MoM_0_ consistently underestimated PVE_GREX_ as a result of ignoring estimation uncertainty.

We also compared IGREX with an existing method in the literature, RhoGE [23]. RhoGE is an LDSC-based approach for estimating PVE_GREX_. However, this method does not adjust for estimation uncertainty. The results are shown in Fig. 1e. As expected, IGREX yielded unbiased estimation while RhoGE substantially underestimated PVE_GREX_ in most settings. It achieved similar accuracy as IGREX only when the genetically regulated expression accounted for most of the expression variation, PVE_*y*_ ≥ 0.9. In other words, RhoGE only works well when the genetically-predicted expression levels are very close to the true underlying expression levels for most of the genes, which may not be realistic for real data analysis.

### Real data applications with individual-level GWAS data

With eQTL data of 48 human tissues from the GTEx project as reference, we applied IGREX to two individual-level GWAS datasets, the Northern Finland Birth Cohorts program 1966 (NFBC) [32] and the Wellcome Trust Case Control Consortium (WTCCC) [33]. The details of the datasets and the data pre-processing procedures are described in the Methods Section.

In analyzing the NFBC data, we focused on six quantitative traits with statistically significant heritability, based on 5, 123 individuals and 309, 245 genotyped SNPs. Those six traits are Glucose 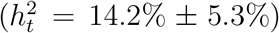, high-density lipoprotein cholesterol (HDL, 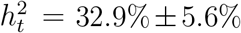), low-density lipoprotein cholesterol (LDL, 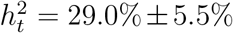), triglycerides (TG, 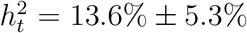), total cholesterol (TC, 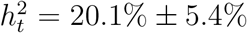) and systolic blood pressure (SysBP, 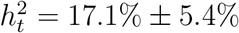). Fig. 2a-b shows the tissue-specific 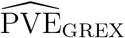 estimates of the six traits. The REML and MoM methods yielded similar estimates in most of the tissues.

**Figure 2:**
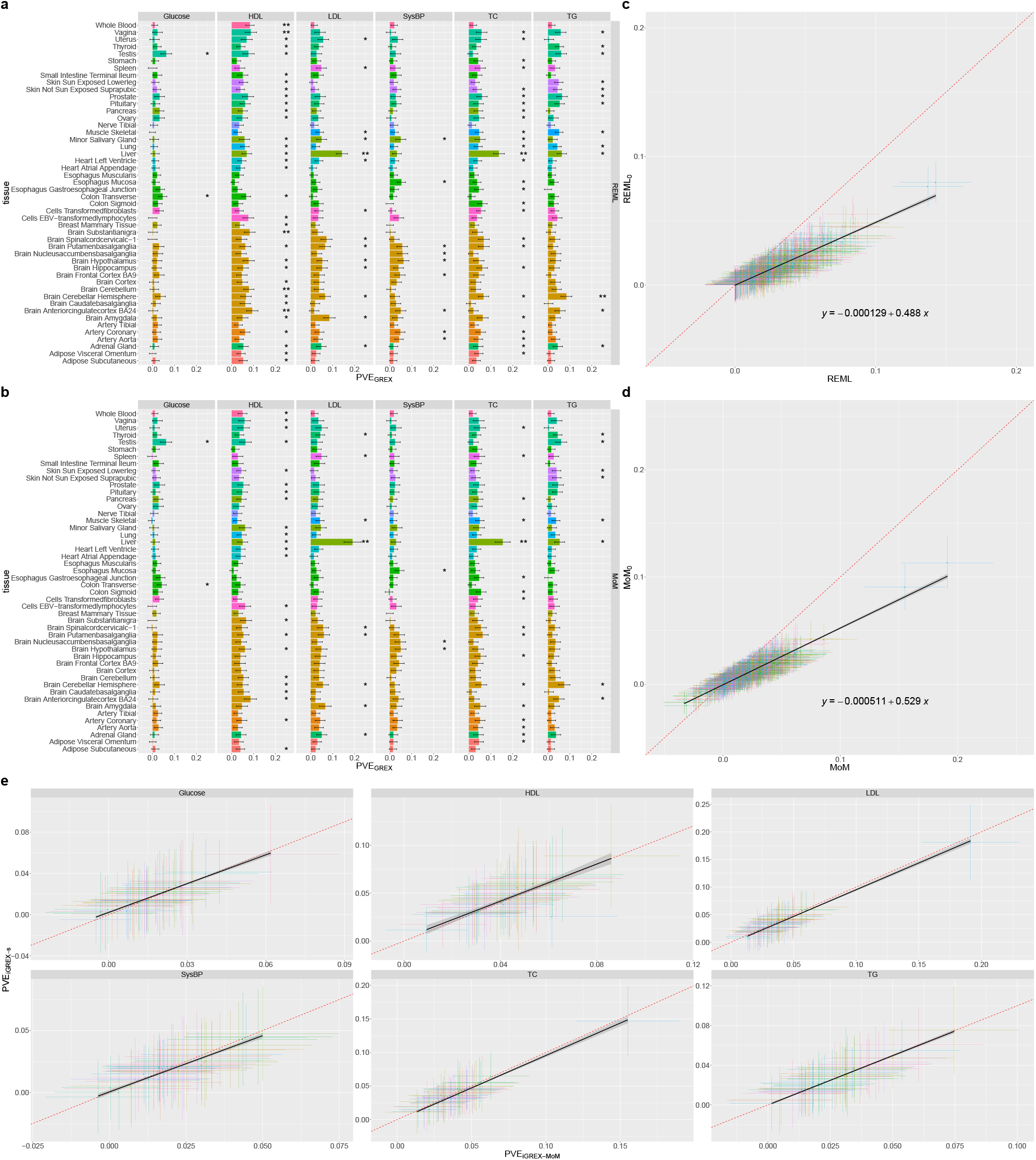
Tissue-specific 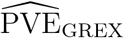 of the six traits from NFBC data set. (**a-b**) 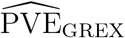 obtained by REML and MoM. Tissues are colored according to their categories. The number of asterisks represents the significance level: *p*-value< 0.05 is annotated by *; *p*-value< 0.05/48 is annotated by **. (**c-d**) All pairs of estimates generated by REML and MoM against their counterparts without accounting for uncertainty. A regression line is fitted and the estimated coefficients are given in the plot. (**e**) Each panel is a plot of 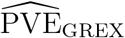 generated by IGREX-s against those generated by MoM for all 48 tissues in one of the six traits.

IGREX can also be used to inform trait-relevant tissue types. By testing *H*_0_: PVE_GREX_ = 0 in each tissue type, we observed significant GREX components in liver for both LDL and TC. As shown in Fig. 2a, 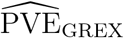 for LDL in liver is as high as 14.3% (with standard error 2.6%), capturing 52.6% of total heritability defined as 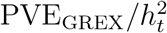; and TC also has 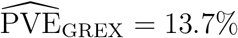 (with standard error 2.5%) in liver, which captures 79.4% of total heritability (see Supplementary Fig. 6). It is known that LDL synthesized in liver is an important lipoprotein particle for transporting cholesterol in the blood [34, 35]. Our findings suggest that genetic variants affect LDL through regulating their corresponding gene targets and liver is the most relevant tissue involved in gene regulation. Next, we analyzed the impact of ignoring the estimation uncertainty (with the complete results given in the Supplementary Fig. 5). As shown in Fig. 2c-d, the 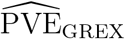 declined substantially as a result of ignoring expression estimation uncertainty. In Fig. 2e, we compared the estimates based on individual level data using IGREX-i versus those based on IGREX-s with summary statistics. For all six of the traits, the IGREX-s estimates well approximated the estimates using the individual level data, which is consistent with our simulation results.

Next we investigated the role of GREX in complex human traits and diseases, using the WTCCC dataset [33]. We applied IGREX to estimate the tissue-specific PVE_GREX_ of seven diseases including bipolar disorder (BD), coronary artery disease (CAD), Crohn’s disease (CD), hypertension (HT), rheumatoid arthritis (RA), type 1 diabetes (T1D) and type 2 diabetes (T2D). The estimates of 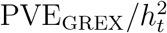 obtained by REML are shown in Supplementary Fig. 8. The top GREX components measured by 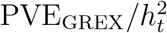 are 12.8% for BD in amygdala, 21.2% for CAD in spinal cord, 18.4% for CD in amygdala, 16.7% for HT in spleen and 17.9% for T2D in anterior cingulate cortex. The average estimates of 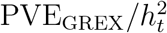 across 48 tissues for RA and T1D are as high as 34.1% and 71.2%, respectively. Both RA and T1D are autoimmune diseases, with well-established strong associations in the major histocompatibility complex (MHC) region [33, 36]. After removing the MHC region, we observed a substantial reduction in the 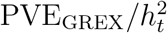 estimates: the mean 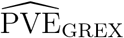 dropped from 34.1% to 7.6% for RA and from 71.2% to 11.7% for T1D, as shown in Fig. 3a. Additionally, the tissue-specific comparisons presented in Fig. 3b showed an extensive reduction of PVE_GREX_ in all tissue types for T1D and RA, while such changes were not observed for other traits. This finding suggests the heavy involvement of GREX variation in the immune functions related to the MHC region for both RA and T1D. Here we illustrate that IGREX can be used to inform disease/trait-relevant tissue types or cellular contexts.

**Figure 3:**
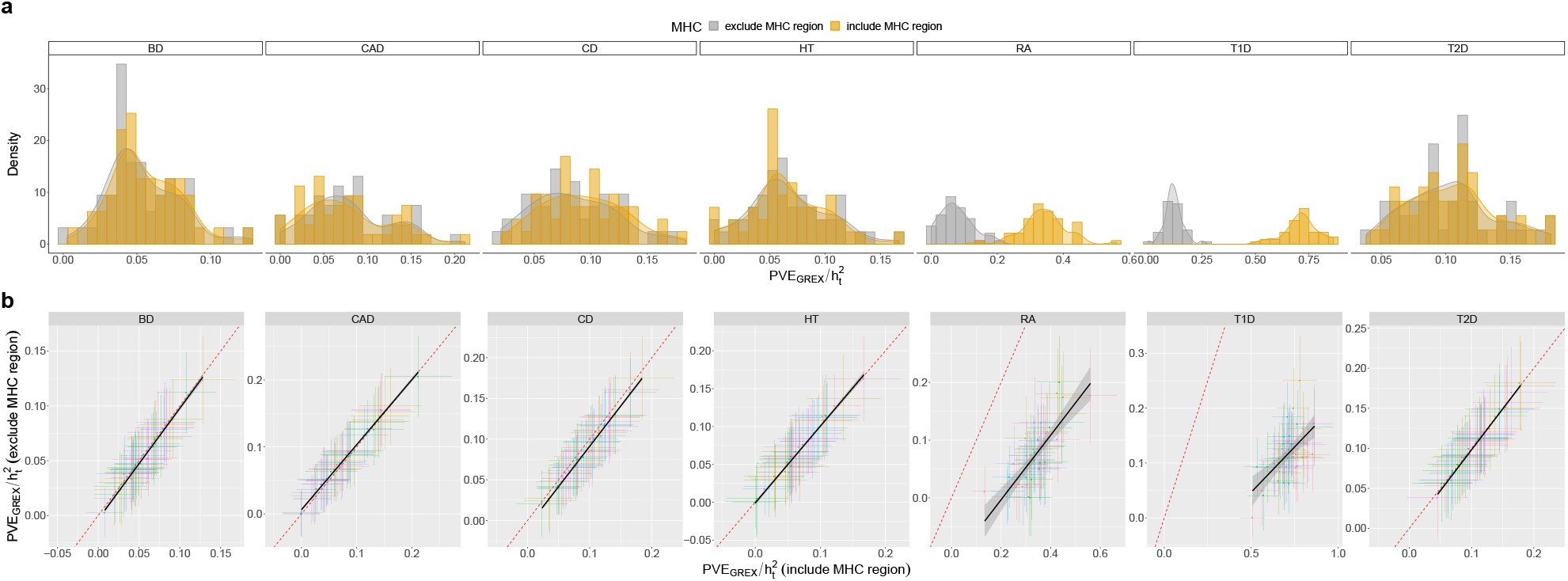
Percentage of heritability explained by GREX 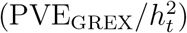 of the seven traits from WTCCC data. (**a**) The distributions of estimated 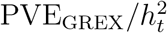 across 48 GTEx tissues. (**b**) Tissue-specific comparisons of 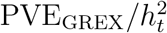 estimated by whole genome with those estimated by excluding the MHC region.

### Analysis of a wide spectrum of phenotypes using IGREX-s with summary-level

**GWAS data** The vast amount of publicly available summary-level GWAS data and their easy accessibility allow us to conduct a comprehensive evaluation of the impact of GREX on a wide spectrum of phenotypes using IGREX-s, from molecular traits such as proteins and metabolites to various complex phenotypes including schizophrenia, height, and body mass index (BMI). In the following analysis, we used the genotypes of the 635 GTEx samples as the LD reference 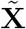 in the IGREX-s estimation.

First, we estimated PVE_GREX_ in 249 proteins with significantly nonzero heritabilities using summary statistics from a plasma protein quantitative trait loci (pQTL) study [38], as summarized in Fig. 4a. In Supplementary Fig. 10, the heritabilities estimated by IGREX 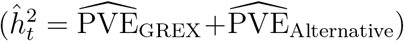 are shown to be highly consistent with those estimates obtained using MoM [37]. From this perspective, heritability can be attributed to two components: the GREX component and its alternative effects. Then, we grouped 48 tissue types into 16 groups by their functions and tested the significance of tissue-specific GREX effects on the 249 proteins. We observed a significant GREX contribution in many tissue-protein pairs (Fig. 4b and Supplementary Fig. 11-13). In particular, 9 out of the 249 proteins had significant GREX components in at least one tissue type at 0.05 level after Bonferroni correction. As shown in Fig. 4d-e, some proteins, including CD96, DEFB119, MICB and PDE4D, exhibit cross-tissue GREX impacts; meanwhile other proteins, namely CFB, CXCL11, EVI2B, IDUA and LRPAP1, have tissue-specific GREX effect patterns. We found these tissue-specific patterns to be consistent with protein functions. For example, the CFB protein, which is implicated in the growth of preactivated B-lymphocytes, is found to be most associated with GREX in EBV-transformed lymphocytes 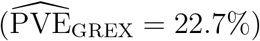. As another example, the CXCL11 protein has the highest 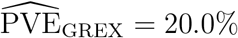 in pancreas, and the *CXCL11* gene is often over-expressed in pancreas tissue [39]. We also noted that 6 out of the 9 proteins were immune-related, echoing our previous implications of the important role of GREX in immune processes. In addition to the proteins, metabolic traits are also important intermediate traits for complex biological processes. We applied IGREX-s to a summary level data set of circulating metabolites [40], and studied the impact of GREX on metabolic traits. The results are discussed in the Supplementary Materials.

**Figure 4:**
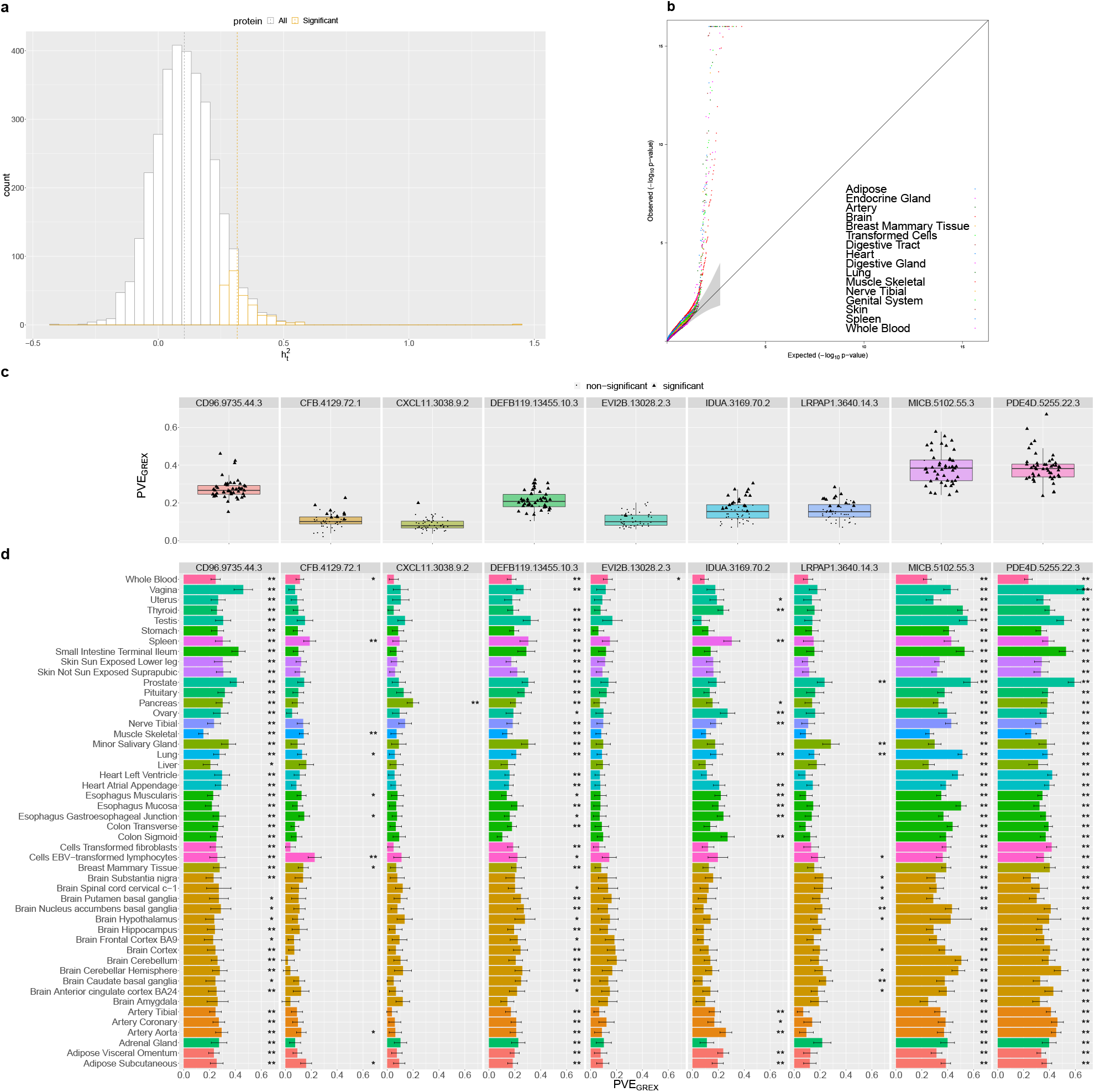
Analysis of plasma pQTL summary statistics. (**a**) The distribution of estimated heritabilities of 3, 283 proteins estimated using [37]. The whole study is colored in grey, while the 249 proteins with significant heritabilities are colored in yellow. Dashed lines represent the means of corresponding distributions. (**b**) QQ-plot of PVE_GREX_ *p*-values of tissue-protein pairs. GTEx tissues are categorized into 16 types and colored accordingly. (**c**) The Manhattan plot of the protein encoding genes in aorta, cerebellum, liver and whole blood. Each point represents a tissue-protein pair. (**d**) 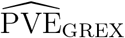 in the 9 proteins whose 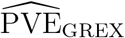 are significant in at leat one tissue at 0.05 level using Bonferrni correction. (**e**) 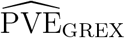 obtained by IGREX-s. Tissues are colored according to their categories. The number of asterisks represents the significance level: *p*-value< 0.05/48 is annotated by *; *p*-value< 0.05/(48 * 9) is annotated by **.

Then we applied IGREX-s to the summary data of complex human traits. Here we analyzed three traits: schizophrenia (SCZ), height, and BMI. We considered four datasets of schizophrenia with increasing and overlapping samples: SCZ subset [41], SCZ1 [42], SCZ1+Sweden (SCZ1Swe)[43] and SCZ2 [44]. We found that the estimated 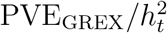 in all four SCZ datasets have higher values in the brain tissues than in other tissue types (Fig. 5b). As expected, the statistical power increases with sample size of GWAS (Fig. 5a). Additionally, we also analyzed the human height and BMI phenotypes using pairs of independent GWAS data for replication purposes. The obtained estimates, 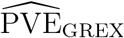, from pairs of independent GWAS data are highly consistent. Although the analysis results are reproducible in several different data sets, we noted the estimated percentages of heritability explained by GREX for all three complex traits are less than 10% (8.7% for schizophrenia, 8.7% for height and 3.7% for BMI in the most expressed tissue types. See Fig. 5c and Supplementary Fig. 15).

**Figure 5:**
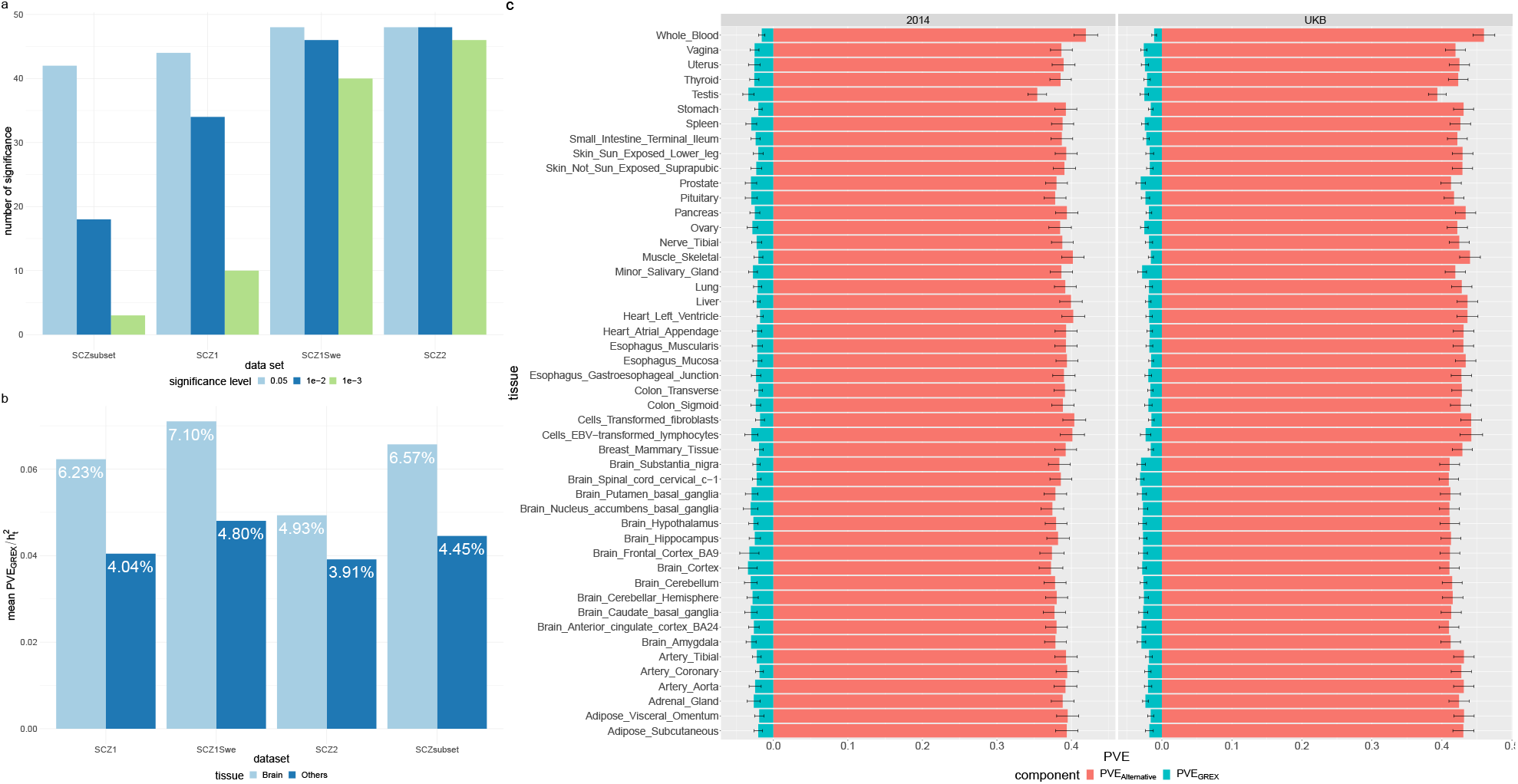
Analyses of complex traits: schizophrenia and height. (**a**) Number of significant GREX components revealed under different significance levels for the four schizophrenia datasets. (**b**) Mean estimated percentages of heritability for schizophrenia explained by GREX in brain tissues and in other tissues. (**c**) 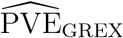 and 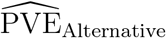 of height estimated using height2014 and UKB datasets, respectively.

The relatively low GREX contribution to complex traits other than lipid or molecular traits can be attributed to multiple reasons. First, it is known that trans-acting genetic effects can explain a substantial proportion of expression variation [8, 12]. However, trans-eQTL effects are often tissue-specific and can be harder to detect and replicate across studies [45]. In TWAS-types of analysis, generally the prediction of gene expression is based on only cis genetic variants of each gene. As such, the PVE_GREX_ values reported here, also based on only cis genetic variants, may be underestimated. In the next section, we will further explore the contribution of trans-eQTLs. Second, the genetic effects on gene expression may not be steady across the reference GTEx data with largely non-diseased tissues for general purposes and the GWAS data with diseased individuals from specific populations [46]. From this perspective, before analyzing specific complex traits and diseases via TWAS, it would be helpful to first estimate the impact of GREX and select the most informative available eQTL reference data.

### Additional insights on GREX considering trans-eQTLs and genetically-regulated alternative splicing events

The cis-acting genetic effects on local gene expression levels are often shared across tissue types and are often replicable across studies [47]. It is also reported that a substantial proportion (up to 70%) of gene expression heritability can be attributed to trans-acting genetic effects which act predominantly in a tissue-specific manner and have a lower rate of replication across studies [48, 49]. More recently, the eQTLGen consortium [15] has conducted a blood-eQTL meta-analysis and has reported 6,298 (31%) trans-eQTL genes for 10,317 trait-associated SNPs using 31,684 blood samples from 37 datasets. The results suggest that trans-eQTLs are prevalent in the genome, while it is still underpowered to detect them for tissues other than whole blood given the often tissue-specific nature of trans-genetic effects and the limited sample sizes for most tissue types.

Although it is still unrealistic to account for all trans-eQTLs in the estimation of PVE_GREX_ due to the limitation of sample sizes, it is possible to explore the potential by incorporating the blood-based trans-eQTLs reported by the eQTLGen consortium and re-estimating 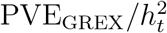. We first analyzed 13 datasets comprised of 12 phenotypes that have significant 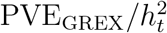 estimates in the whole blood, including 7 proteins, 1 lipid trait and 4 complex diseases (with 2 SCZ datasets). We observed an increasing trend of 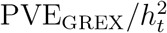 in the blood for all 13 datasets (Fig. 6a), by accounting for only ~ 1, 700 unique trans-eQTLs that are not cis-eQTLs. As a comparison, we applied the same procedure to 13 GTEx brain tissues of the two largest SCZ datasets, and did not observe an increase in 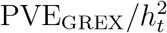 (Fig. 6b). This is not surprising because the trans-eQTLs incorporated above were detected and reported based on whole blood samples and may not be trans-eQTLs in the brain tissues. Our results suggest that the estimation of GREX impacts on traits can be further boosted by incorporating robust trans-eQTLs from the same tissue types.

**Figure 6:**
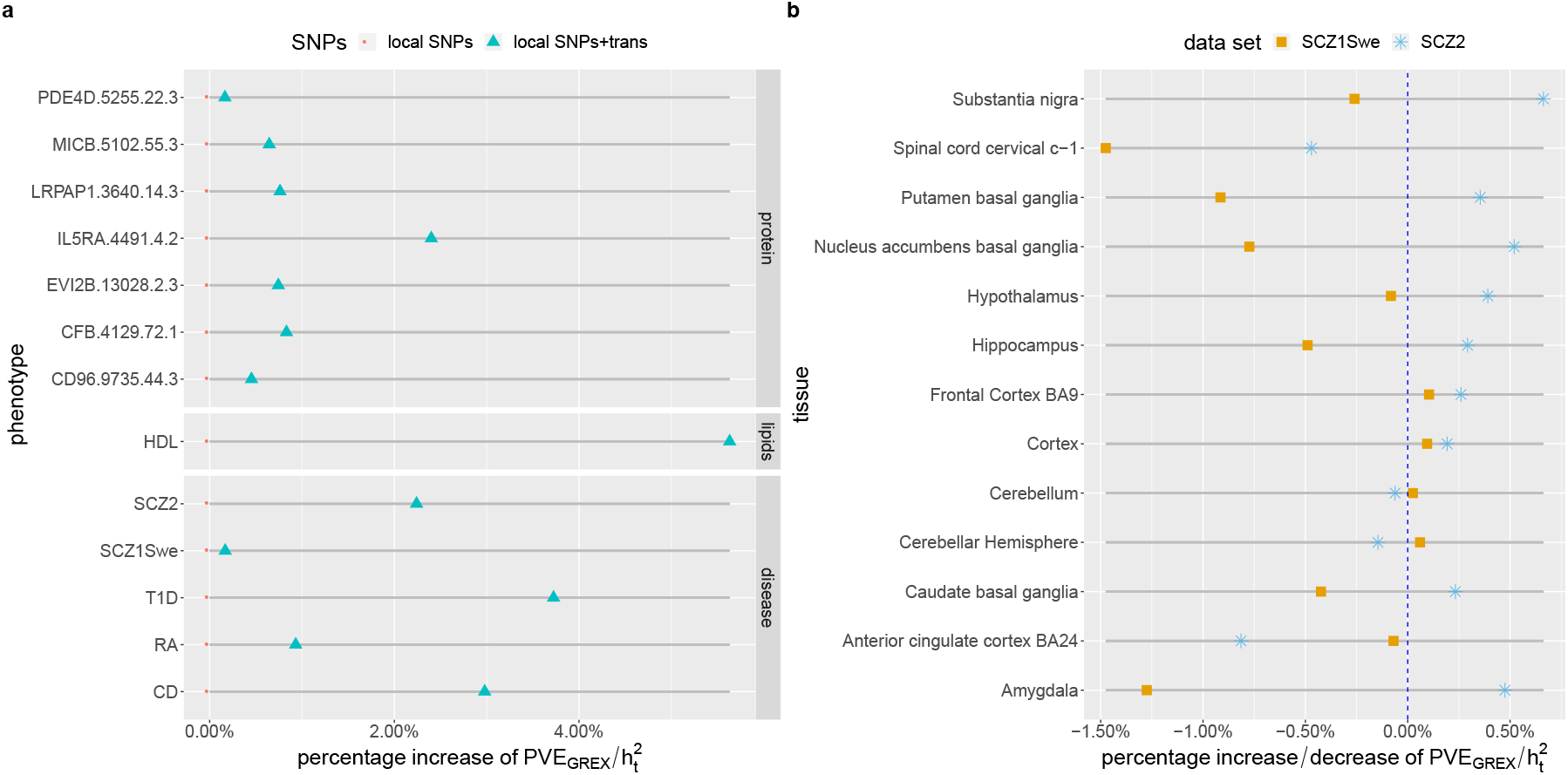
Comparison of 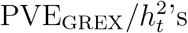 estimated with only local SNPs and those estimated with additional trans-eQTLs. (**a**) Estimated 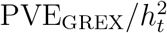 of 13 datasets in blood. All these datasets have significant 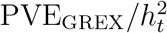 in blood at 0.05 nominal level using only local SNPs. (**b**) Estimated 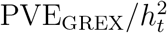 of two largest SCZ data sets in 13 GTEx brain tissues. All these tissues have significant 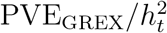 at 0.05 nominal level in both datasets using only local SNPs.

In addition to the gene expression level, we also evaluated the effects of alternative splicing on complex trait heritability. We applied IGREX to quantify the impact of genetically regulated alternative splicing on multiple phenotypes. Alternative splicing is an important gene regulatory process that results in multiple transcripts from a single multi-exon gene. It is commonly observed in humans and plays an essential role in cellular differentiation [50, 51]. Differential variations in splicing may also result in phenotypic variation and contribute to the development of complex diseases including cancer [52, 53, 54]. In a recent work, by extending the TWAS framework to analyze splicing events and associating 40 complex traits with genetically-predicted splicing quantification, novel putative disease-associated genes were detected [55]. Here, using multi-tissue splicing quantification data from GTEX as reference, we applied IGREX to study the impact of genetically-regulated splicing events on four trait-tissue pairs that were found to have a high 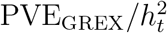. We estimated the proportion of phenotypic variation explained by genetically-regulated splicing to be 12.5%, 13.5%, 1.0% and 1.1% for LDL in liver, TC in liver, SCZ in amygdala and SCZ in cerebellar hemisphere, respectively. Unlike eQTLs that are often found to be near transcription starting sites, most of the sQTLs were found to be enriched within gene bodies, in particular within the introns they regulate, and have little to no effects on cis gene expression levels [55, 56]. In other words, sQTLs are often independent of eQTLs. Therefore, integrating genetically-regulated splicing quantification may partially explain the phenotypic variation attributed to alternative genetic factors, PVE_Alternative_. We argue that with the proper multi-omics reference data, similar analyses can be conducted to quantify the impact of genetically-regulated methylation, protein, and other multi-omics variation on phenotype [51].

## Discussion

In this work, we proposed a method, IGREX, for integrating GWAS and eQTL reference data to quantify the GREX impact on phenotype. IGREX can be applied to both individual-level and summary-level GWAS data, and was shown to achieve estimation accuracy even when the eQTL effects are weak. IGREX can be used in many ways: it can inform the role of GREX variation in various phenotypes and/or the role of GREX in known pathways; it can guide the selection of eQTL reference data and suggest trait-relevant tissues/cell-types/contexts; and it is generally applicable to the integration of GWAS with other omics data types to examine the role of genetically-regulated multi-omics traits.

IGREX is closely related to several existing methods and here we briefly discuss the connections and distinctions. By also integrating an eQTL reference and GWAS data, methods including TWAS [17], PrediXcan [16], and the more general MetaXcan [21] aim to identify specific trait-associated genes. In contrast, IGREX estimates the impact of genetically regulated expression from a global perspective by quantifying the phenotypic variation that can be attributed to the GREX component. Since both the TWAS-type of analyses and IGREX rely on the shared GREX variation across eQTL and GWAS data, we argue that with the increasing availability of eQTL resources in different populations, conditions and contexts, the proper selection of eQTL reference panels via IGREX will greatly promote the chances of successes in the subsequent TWAS-type of analyses.

There are also existing methods, such as RhoGE, designed for identifying and estimating correlations between gene expression and complex traits. RhoGe provides an LDSC-based approach for estimating PVE_GREX_. Unlike IGREX, this method does not adjust for estimation uncertainty. Consequently, it significantly underestimates the PVE_GREX_ when the eQTL effects on expression levels are weak or moderate. In fact, RhoGE estimated the PVE_GREX_ for the majority of 1, 350 tissue-trait pairs to be almost negligible, with the first quantile, the median, and the third quantile being 0.00125%, 0.162% and 0.616%, respectively [23]. In contrast, as demonstrated via simulation studies, IGREX can accurately estimate PVE_GREX_ in various scenarios by accounting for the estimation uncertainty.

Based on estimating PVE_GREX_ for a wide-array of tissue-trait pairs, we observed a stronger impact of GREX on molecular intermediate traits and lipid traits in trait-relevant tissue types. We also observed a relatively low PVE_GREX_ for complex traits in general. The big picture suggests the attenuated impact on downstream phenotypes (e.g, height and SCZ), which is consistent with the result from a pioneer study [57]. However, we noted that the PVE_GREX_ estimates could be improved. A substantial amount of expression heritability is explained by trans-acting genetic factors while current TWAS and IGREX analyses are mainly using only cis-eQTLs. We explored the potential of incorporating trans-eQTLs in TWAS analysis by re-estimating PVE_GREX_ for selected traits in blood tissues with significant trans-eQTLs independently derived from the blood-based eQTLGen Consortium. We observed consistent increases in PVE_GREX_ for blood-related traits. In contrast, such an increase was not observed in the PVE_GREX_ estimates for other tissue types, again illustrating the importance of considering trait-relevant tissue types/conditions in the TWAS-type of analyses. Additionally, we extended the IGREX analysis to quantify the impact of genetically-regulated alternative splicing events on selected traits. Our results suggested the potential for extending TWAS-type of analysis to integrate reference multi-omics QTL data with GWAS in mapping novel disease/trait-associated genes with mechanisms via other omics traits (such as splicing, methylation, protein, etc.).

A key assumption in applying IGREX or TWAS methods with a general-purpose eQTL data as reference is the existence of steady-sate component in GREX, i.e., the genetic effects on gene expression *β_g_* are shared across the eQTL reference and GWAS data. However, there are many situations in which this assumption is violated. For example, it has been observed that CAD-risk SNPs have a larger overlap with cis-eQTLs isolated from disease-relevant tissues than those from GTEx tissues [46], implying the existence of a dynamic component. In the presence of this dynamic component, the accuracy of 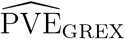 based on GTEx is reduced. In those cases, we suggest exploring other trait-relevant or condition-specific eQTL reference panels using IGREX for a better understanding of the role of GREX and before conducting TWAS analysis.

## Methods

### The IGREX-i for individual-level GWAS data

First, let 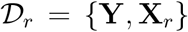 denote the reference data set from an eQTL study, where **Y** ∈ ℝ^*n_r_* × *G*^ is the gene expression matrix, **X**_*r*_ ∈ ℝ^*n*_*r*_ ×*M*^ is the genotype matrix, *n_r_* is the sample size of the eQTL study, *G* is the number of genes and *M* is the number of single-nucleotide polymorphisms (SNPs). Suppose we have individual-level GWAS data 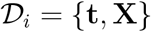 comprised of phenotype vector **t** ∈ ℝ^*n*^ and genotype matrix **X** ∈ ℝ^*n* × *M*^, where *n* is the GWAS sample size. For *g* = 1,…, *G*, we let the *g*-th gene expression vector **y**_*g*_ ∈ ℝ^*n_r_*^ denote the corresponding column of **Y**, local genotype matrices **X**_*r,g*_ ∈ ℝ^n_*r*_ ×*M_g_*^ and **X**_*g*_ ∈ ℝ^*n*×*M_g_*^ denote the corresponding *M_g_* columns in **X**_*r*_ and **X**, respectively, where *M_g_* is the number of local SNPs for *g*-th gene. To make the notation uncluttered, we further assume that **X**_*r,g*_ and **X**_*g*_ have been standardized and both **y**_*g*_ and **t** have been properly adjusted for covariates. The complete model that accounts for covariates is described in the Supplementary Materials. Now, we consider linear model (1) that associates the gene expression vector **y**_*g*_ to **X**_*r,g*_:

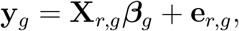

where ***β**_g_* is an *M_g_* × 1 vector of genetic effects on the gene expression levels, 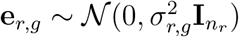 is a vector of independent noise and **I** is the identity matrix with the subscript being its size. Assuming that there is a steady-state component in gene expression regulated by genetic variants, individuals in 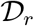 and 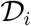 share the same ***β**_g_*. Hence, the GREX in 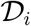 can be evaluated by **X**_*g*_***β***_*g*_. Then, we assume that the phenotype **t** can be decomposed into two parts, i.e., the genetic effects via GREX and the genetic effects through alternative pathways, as in model (2):

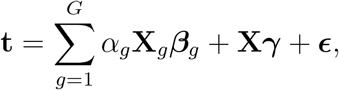

where *α_g_* is the effect of **X**_*g*_***β***_*g*_ on **t**, *γ* is an *n* × 1 vector of alternative genetic effects and 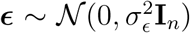 is a vector of independent errors. The term 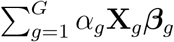 can be viewed as the overall impact of GREX on the phenotype and **X**_***γ***_ represents the alternative impact of genotypes on the phenotype. Given a genotype vector **x** ∈ ℝ^*M*^ and a phenotype *t* ∈ ℝ, the impact of GREX can be quantified by the proportion of variance explained by the GREX component:

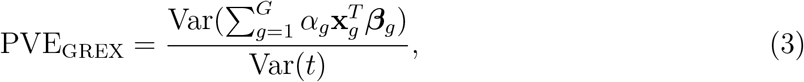

where **x**_*g*_ is the subvector of genotype **x** corresponding to the *g*-th gene.

To estimate PVE_GREX_, we introduce the following probabilistic structure for the effects in model (1) and (2):

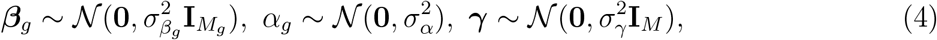

which is motivated by a recent theoretical justification [58] for heritability estimation on a mis-specified linear mixed model (LMM). This prior specification in (4) provides a great computational advantage as well as a stable performance for IGREX under model mis-specification, as demonstrated in the simulation studies.

The proposed method for individual-level GWAS data, IGREX-i, provides a two-stage framework for estimating PVE_GREX_. In the first stage, we estimate the parameters 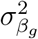 and 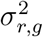 in model (1) via a fast expectation-maximization (EM)-type algorithm, the parameter-expanded EM (PX-EM) algorithm [59]. Based on the estimates, denoted as 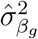 and 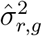, the posterior distribution of ***β**_g_* is given by

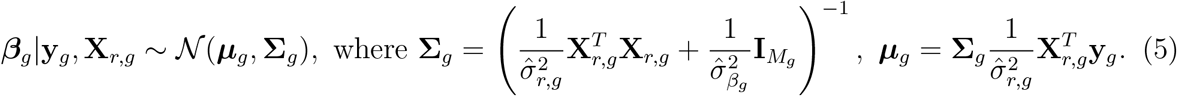

In the second stage, we treat the posterior distribution obtained in (5) as the prior distribution of ***β**_g_* in model (2). This substitution naturally accounts for the uncertainty in estimating ***β**_g_* which has been captured by **Σ**_*g*_. To evaluate the covariance of **t**, we first note that 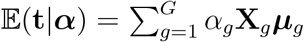 and 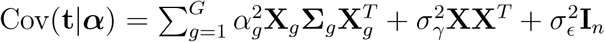; then, using the law of total expectation and total variance, we obtain 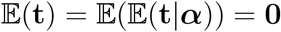 and

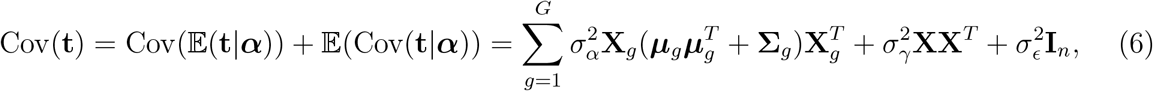

respectively. By observing the form of (6), it is clear that the *i*-th diagonal element of 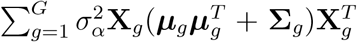 and 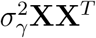 represents the variance explained by GREX and alternative genetic effects, respectively. Therefore, the PVE_GREX_ defined in (3) can be estimated by

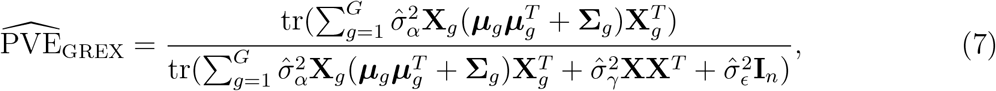

where 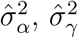 and 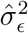 are the estimated values of 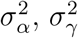 and 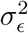, respectively.

IGREX-i provides two methods for estimating the parameters and 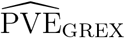 in the second stage. Let 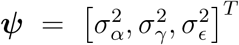 be the vector of parameters to be estimated, 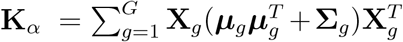 and **K**_*γ*_ = **XX**^*T*^. The first method is based on MoM, which minimizes the distance between the second moment of **t** at the population level and that at the sample level 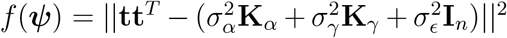. By setting 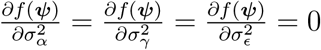, we obtain the estimating equation

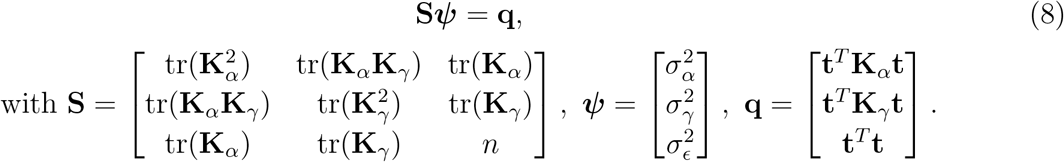

The solution of Equation (8) is given by 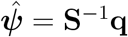. And the variance-covariance matrix of 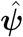 is given by 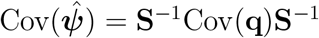 using the sandwich estimator. Then, the standard error of 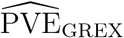 can be obtained by the delta method (see Supplementary Materials). The second method applies the restricted maximum likelihood (REML) by further assuming the normal distribution of **t**: 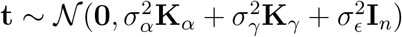. The variance components are estimated by the Minorization-Maximization (MM) algorithm [60].

### The IGREX-s for summary-level GWAS data

The special formulation of method of moments allows IGREX to be extended (IGREX-s) to handle summary-level GWAS data (i.e. *z*-scores) when the individual-level data 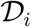 is not accessible. Suppose we only have the *z*-scores from summary-level GWAS data 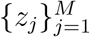 generated from 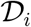. The definition of the *z*-score is 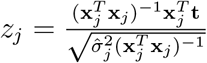, where **x**_*j*_ is the *j*-th column of **X** and 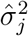 is the estimate of residual variance by regressing **x**_*j*_ on **t**. By assuming that *z*-scores are calculated from a standardized genotype matrix **X**, we have 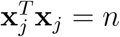. Besides, the polygenicity assumption implies that 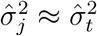, where 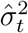 is the estimate of Var(*t*). Hence, we have

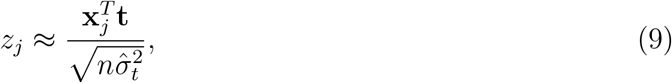

and PVE_GREX_ defined in (3) can be estimated by

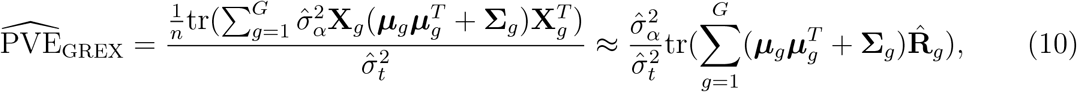

where 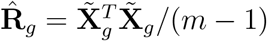 is the estimated LD matrix associated with the *g*-th gene and 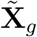 is the corresponding columns of a reference genotype matrix 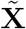. In practice, 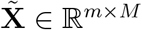 can be the genotype matrix either from the GTEx Project or the 1000 Genomes Project. Now, we consider MoM in the estimating equation (8) to obtain 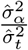. By eliminating 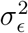 and dividing both sides by *n*^2^, we have

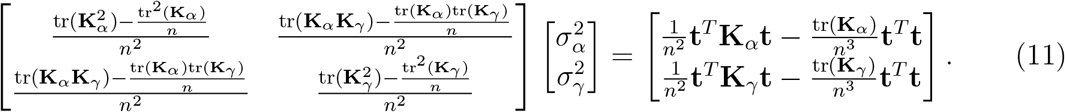

The terms on the left hand side do not involve **t** and thus can be approximated using 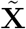 [37]. For example, 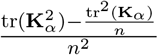 can be well approximated by 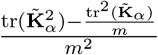, where 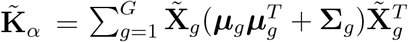. Other terms on the left hand side can be approximated in the same way. For the right hand side, each term can be approximated using 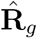 and *z*-scores from approximation (9): 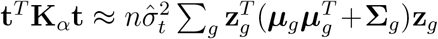, where **z**_*g*_ ∈ ℝ^*M_g_*^ is the vector of *z*-scores corresponding to the *g*-th gene; 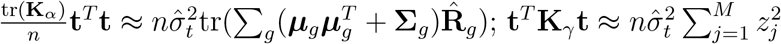; and 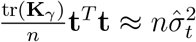. With these approximations, Equation (11) becomes

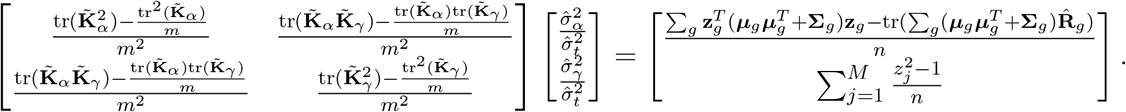

Then, 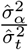 can be obtained by solving this equation. Plugging this estimate into Equation (10) gives the 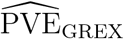. The standard errors of 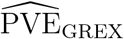 can be estimated by the block jackknife method [61] (Supplementary Materials).

IGREX can incorporate fixed effects to adjust for possible confounding factors, such as population structure. Details are provided in the Supplementary Note.

### GTEx eQTL dataset

We used the gene expression data from the V7 release of GTEx Consortium as our reference dataset. We analyzed the 48 tissues with number of genotyped samples ≥ 70, which are collected from 620 donors with total sample size 10,294. The sample size of each tissue ranges from 80 to 491 (details provided in Supplementary Table 5 4). We set the mappability cutoff at 0.9 to filter gene expression measures with lower quality, leaving 16, 333 – 27, 378 genes to be included in our analysis. Based on the third phase of the International HapMap project phase 3 (HapMap3), 1,189, 556 SNPs were included from the GTEx genotyped data for analysis. For each gene, we included only the SNPs within 500kb of the transcription start and end of each protein coding genes. In real data analysis, we used the covariates provided by the GTEx consortium, including genotype principal components (PCs), Probabilistic Estimation of Expression Residuals (PEER) factors, genotyping platform and sex (as described in https://gtexportal.org/home/documentationPage).

Additionally, the GTEx genotype data was used as an LD reference panel when applying IGREX-s to GWAS summary statistics. In this application, we used top 5 PCs as covariates.

### Individual level GWAS datasets

The NFBC dataset is comprised of 5,402 individuals with ten continuous phenotypes related to cardiovascular disease including Glucose, body mass index (BMI), C-reactive protein (CRP), insulin, high-density lipoprotein cholesterol (HDL), low-density lipoprotein cholesterol (LDL), triglycerides (TG), total cholesterol (TC), diastolic 9 blood pressure (DiaBP) and systolic blood pressure (SysBP). There are 364, 590 genotyped SNPs in this dataset. We first excluded the individuals whose reported sex differed from their sex determined from the X chromosome. We then excluded the SNPs with minor allele frequency less than 1%, with missing values in more than 1% of the individuals or with Hardy-Weinberg equilibrium (HWE) *p*-value below 0.0001. This quality control process yields 5,123 individuals with 319, 147 SNPs in NFBC dataset for our analysis. We evaluated the genetic relatedness matrix (GRM) using the processed genotype data and selected the top 20 PCs as covariates in the study.

The WTCCC dataset contains seven disease phenotypes including bipolar disorder (BD), coronary artery disease (CAD), Crohn’s disease (CD), hypertension (HT), rheumatoid arthritis (RA), type 1 diabetes (T1D) and type 2 diabetes (T2D). It includes ~ 2, 000 cases per phenotype and 3, 004 controls with 490, 032 genotyped SNPs. We first removed the individuals with genotyping rate less than 5%. Then we excluded the SNPs satisfying at least one of the following: minor allele frequency less than 5%; genotypes missing in more than 1% samples; HWE *p*-value is below 0. 001. We also removed the individuals with estimated genetic correlation larger than 2.5%. After quality control, around 4, 700 individuals with 300, 000 SNPs were retained for our analysis (See Supplementary Table 1). Based on the obtained data, we calculated the GRM and extracted top 20 PCs as covariates to be included in our analysis.

### GWAS summary statistics

We analyzed ten summary level GWAS datasets: human plasma pQTL data [38], circulating metabolite data [40], four schizophrenia datasets [41, 42, 43, 44], two independent height datasets [62] and European ancestry of BMI datasets with sample age ≤ 50 separated by men and women [63]. The SNPs with missing information (i.e. chromosome, minor allele, allele frequency) were first removed. Following the practice of LDSC [30], we checked the *χ*^2^ statistic of each SNP and excluded those with extreme values (*χ*^2^ > 80) to prevent the outliers that may unduly affect the results. The detailed information is provided in Supplementary Table 2. After pre-processing, the remaining SNPs were further matched with reference data, and this step is automatically conducted in our IGREX software.

### The eQTLGen summary data

We used the trans-eQTLs in blood provided by the eQTLGen Consortium [15]. The trans-eQTL analysis were restricted to known complex trait-associated SNPs. The significant trans-eQTLs were identified by controlling the FDR at 0.05. There were 5, 4786 gene-SNP pairs composed of 6, 298 genes and 3, 853 SNPs. The remaining pairs after matching with both reference and GWAS datasets are summarized in Supplementary Table 5.

## Data availability

The GTEx gene expression data was downloaded from GTEx Consortium website https://gtexportal.org/home/datasets. The GTEx genotype data can be accessed from dbGAP with accession number phs000424.v7.p2. The HapMap3 genotype data is available at ftp://ftp.ncbi.nlm.nih.gov/hapmap/. The NFBC study was downloaded from dbGAP using accession number phs000276.v1.p1. The WTCCC data was obtained from its consortium website https://www.wtccc.org.uk/info/access_to_data_samples.html. The GWAS summary statistics can be accessed using the links provided in Supplementary Table 2. The eQTLGen data can be downloaded from http://www.eqtlgen.org.

## Software

The R software package IGREX is publicly available on GitHub repository: https://github.com/mxcai/iGREX.

## Acknowledgements

We thank Mr. Kevin J. Gleason for proof-reading the work. This work was supported in part by the National Science Funding of China [61501389]; the Hong Kong Research Grant Council [12316116, 12301417 and 16307818]; The Hong Kong University of Science and Technology [startup grant R9405 and IGN17SC02]; Duke-NUS Medical School WBS [R-913-200-098-263]; Ministry of Education, Singapore. AcRF Tier 2 [MOE2016-T2-2-029, MOE2018-T2-1-046 and MOE2018-T2-2-006]. LSC was independently supported by the National Institutes of Health (NIH) grant R01GM108711. The computational work for this article was (fully or partially) performed on resources of the National Supercomputing Centre, Singapore (https://www.nscc.sg).

