## Supplementary Note for "IGREX for quantifying the impact of genetically regulated expression on phenotypes"

### The supplementary figures and text of "IGREX for quantifying the impact of genetically regulated expression on phenotypes"

Mingxuan Cai[1], Lin S. Chen[2], Jin Liu[3]*& Can Yang[1]*

[1]Department of Mathematics, The Hong Kong University of Science and Technology

[2]Department of Public Health Sciences, The University of Chicago

[3]Center for Quantitative Medicine, Duke-NUS Medical School

### 1 Supplementary Figures

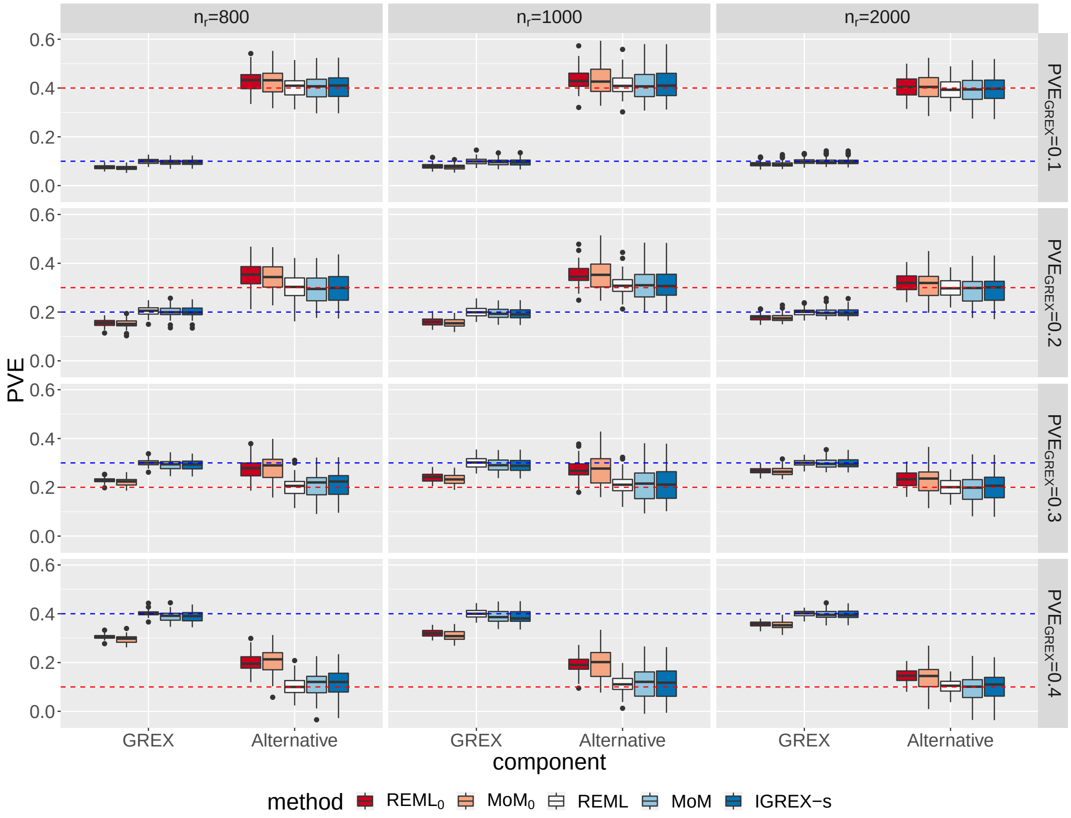

**Supplementary Figure 1:** Estimates of $\text{PVE}_{\text{GREX}}$ and $\text{PVE}_{\text{Alternative}}$ for $\text{REML}_0$, $\text{MoM}_0$, IGREX-REML, IGREX-MoM and IGREX-s with $n = 4000$, $h_t^2 = \text{PVE}_{\text{GREX}} + \text{PVE}_{\text{Alternative}} = 0.5$ and $\text{PVE}_y = 0.3$. $n_r$ is varied at $\{800, 1000, 2000\}$ and $\text{PVE}_{\text{GREX}}$ is varied at $\{0.1, 0.2, 0.3, 0.4\}$. The blue dashed line and red dashed line represent true values of $\text{PVE}_{\text{GREX}}$ and $\text{PVE}_{\text{Alternative}}$, respectively. The results are summarized from 50 replications.

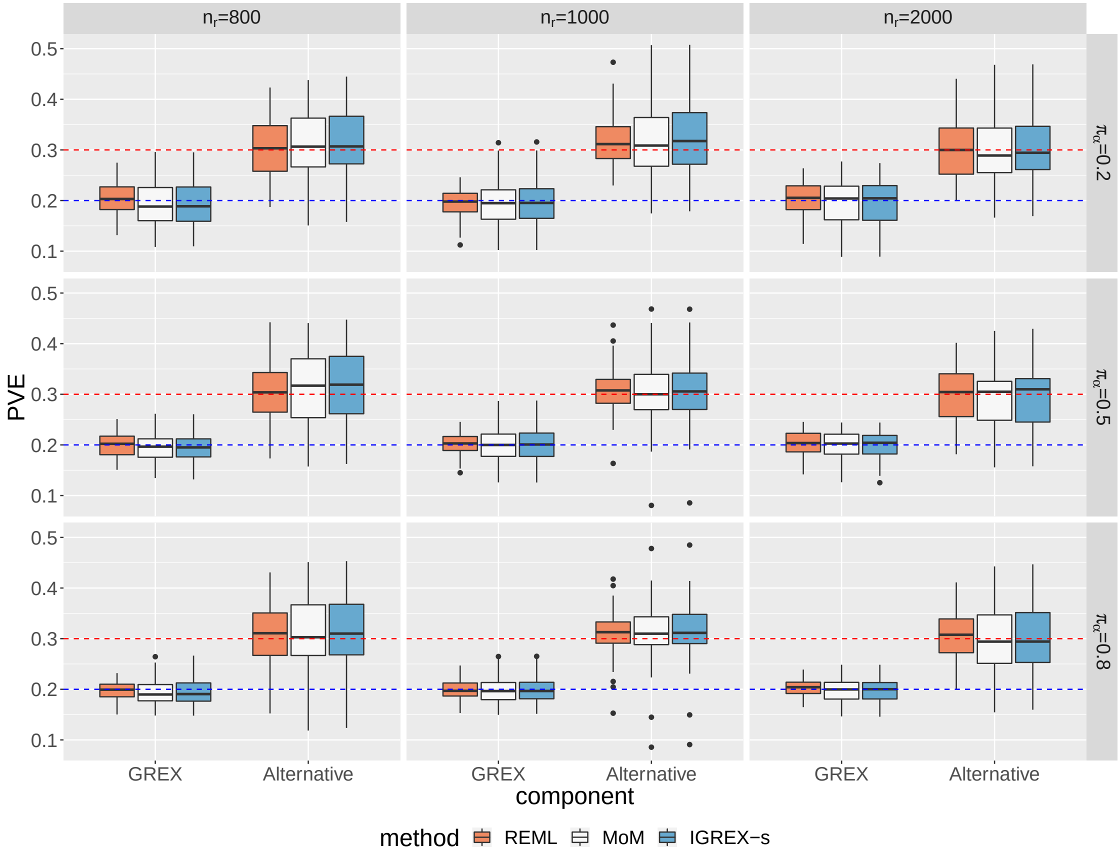

**Supplementary Figure 2:** Estimates of $\text{PVE}_{\text{GREX}}$ and $\text{PVE}_{\text{Alternative}}$ for IGREX-REML, IGREX-MoM and IGREX-s with $n = 4000$, $\text{PVE}_{\text{GREX}} = 0.2$, $\text{PVE}_{\text{Alternative}} = 0.3$, $\text{PVE}_y = 0.3$ and $\pi_\beta = 0.2$. $n_r$ is varied at $\{800, 1000, 2000\}$ and $\pi_\alpha$ is varied at $\{0.2, 0.5, 0.8\}$. The blue dashed line and red dashed line represent true values of $\text{PVE}_{\text{GREX}}$ and $\text{PVE}_{\text{Alternative}}$, respectively. The results are summarized from 50 replications.

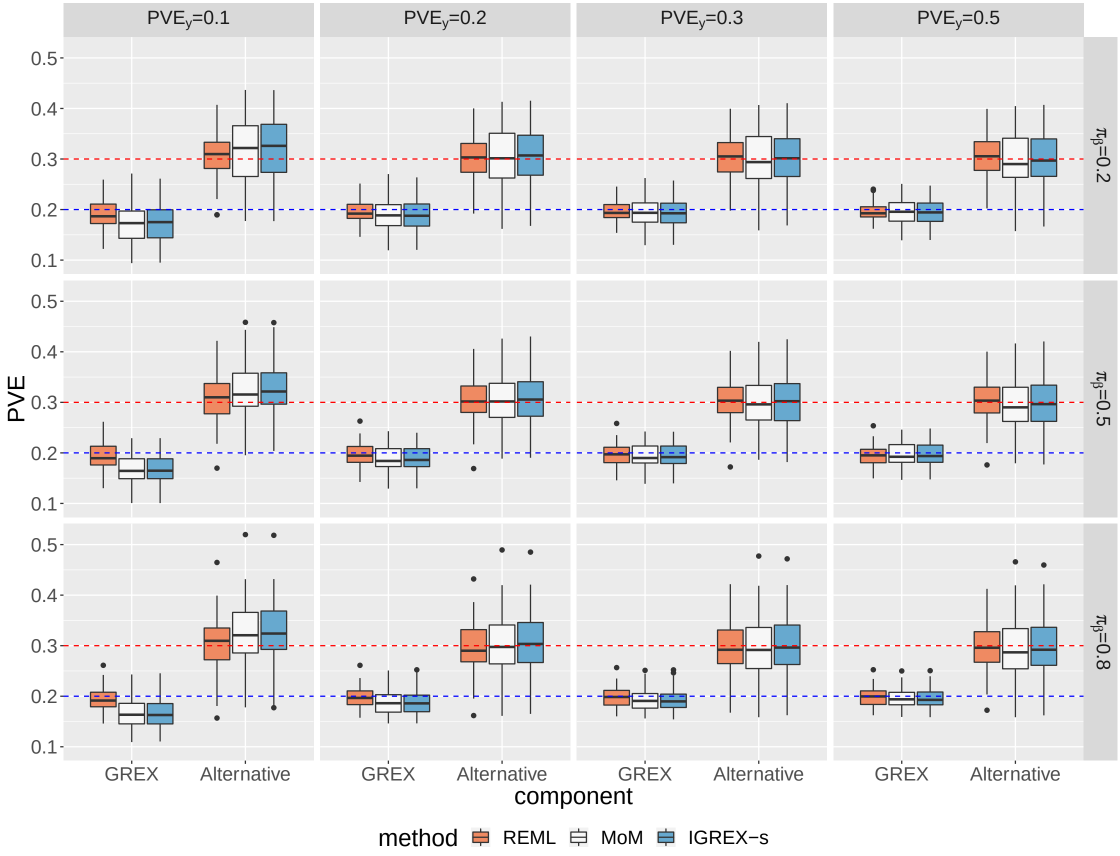

**Supplementary Figure 3:** Estimates of $\mathrm{PVE}_{\mathrm{GREX}}$ and $\mathrm{PVE}_{\mathrm{Alternative}}$ for IGREX-REML, IGREX-MoM and IGREX-s with $n = 4000$, $n_r = 800$, $\mathrm{PVE}_{\mathrm{GREX}} = 0.2$, $\mathrm{PVE}_{\mathrm{Alternative}} = 0.3$ and $\pi_\beta = 0.2$. $\mathrm{PVE}_y$ is varied at $\{0.1, 0.2, 0.3, 0.5\}$ and $\pi_\alpha$ is varied at $\{0.2, 0.5, 0.8\}$. The blue dashed line and red dashed line represent true values of $\mathrm{PVE}_{\mathrm{GREX}}$ and $\mathrm{PVE}_{\mathrm{Alternative}}$, respectively. The results are summarized from 50 replications.

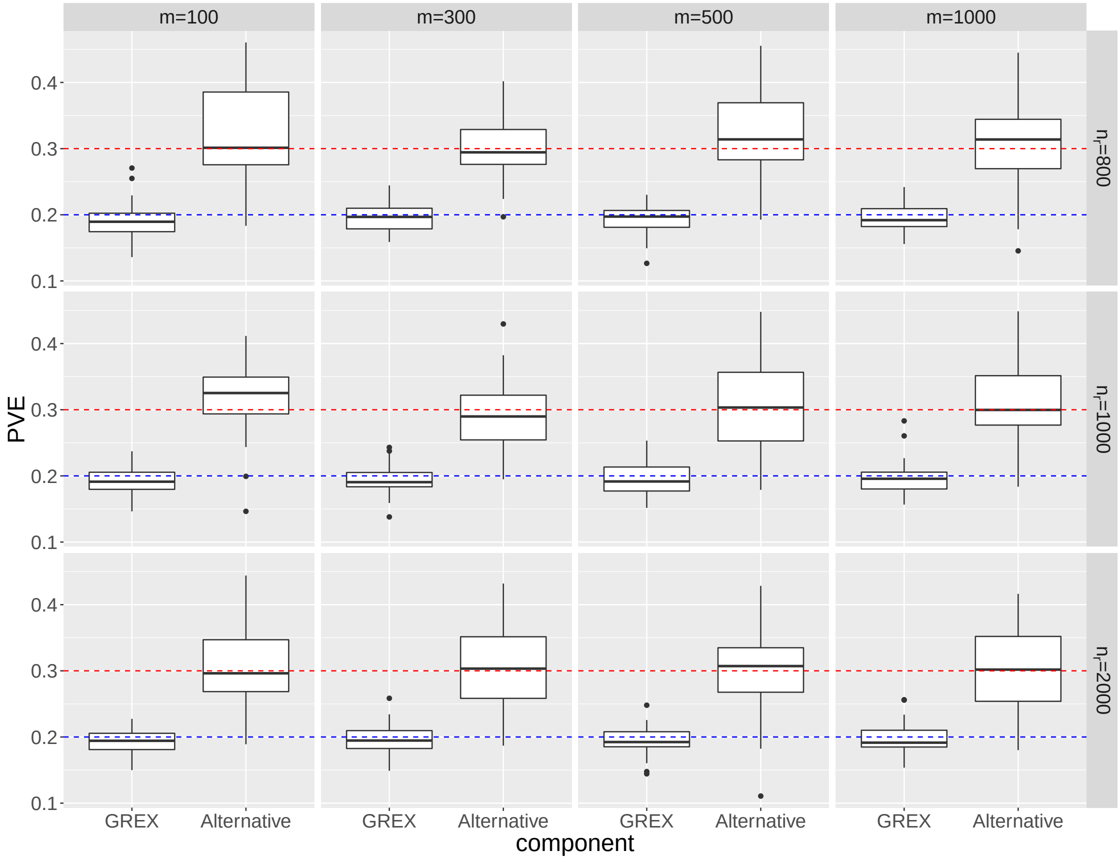

**Supplementary Figure 4:** Estimates of $\text{PVE}_{\text{GREX}}$ and $\text{PVE}_{\text{Alternative}}$ for IGREX-s with $n = 4000$, $\text{PVE}_{\text{GREX}} = 0.2$, $\text{PVE}_{\text{Alternative}} = 0.3$ and $\text{PVE}_y = 0.3$. $n_r$ is varied at $\{800, 1000, 2000\}$ and $m$ is varied at $\{100, 300, 500, 1000\}$. The blue dashed line and red dashed line represent true values of $\text{PVE}_{\text{GREX}}$ and $\text{PVE}_{\text{Alternative}}$, respectively. The results are summarized from 50 replications.

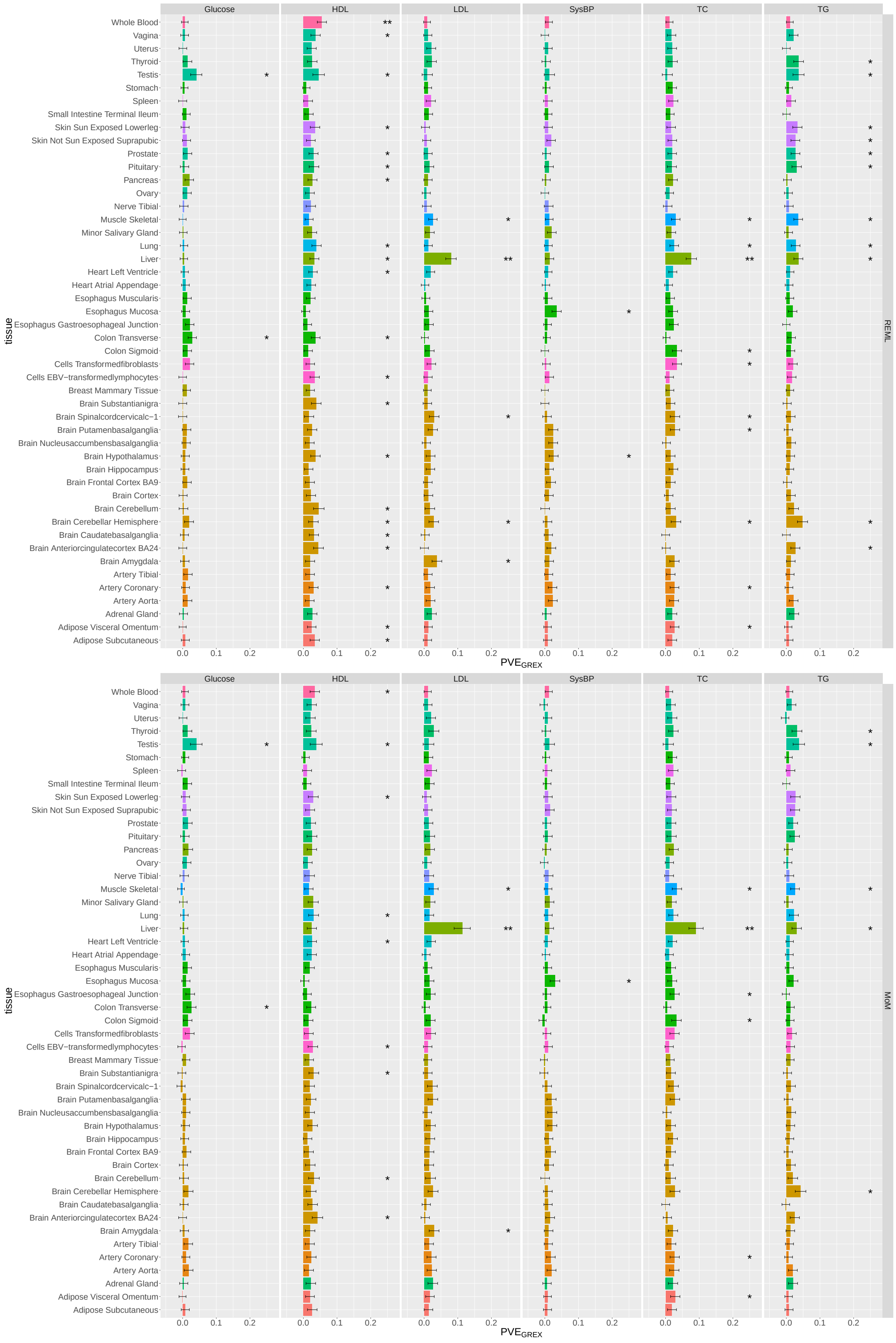

**Supplementary Figure 5:** REML and MoM estimates of PVE$_{\text{GREX}}$ for NFBC data without accounting for uncertainty.

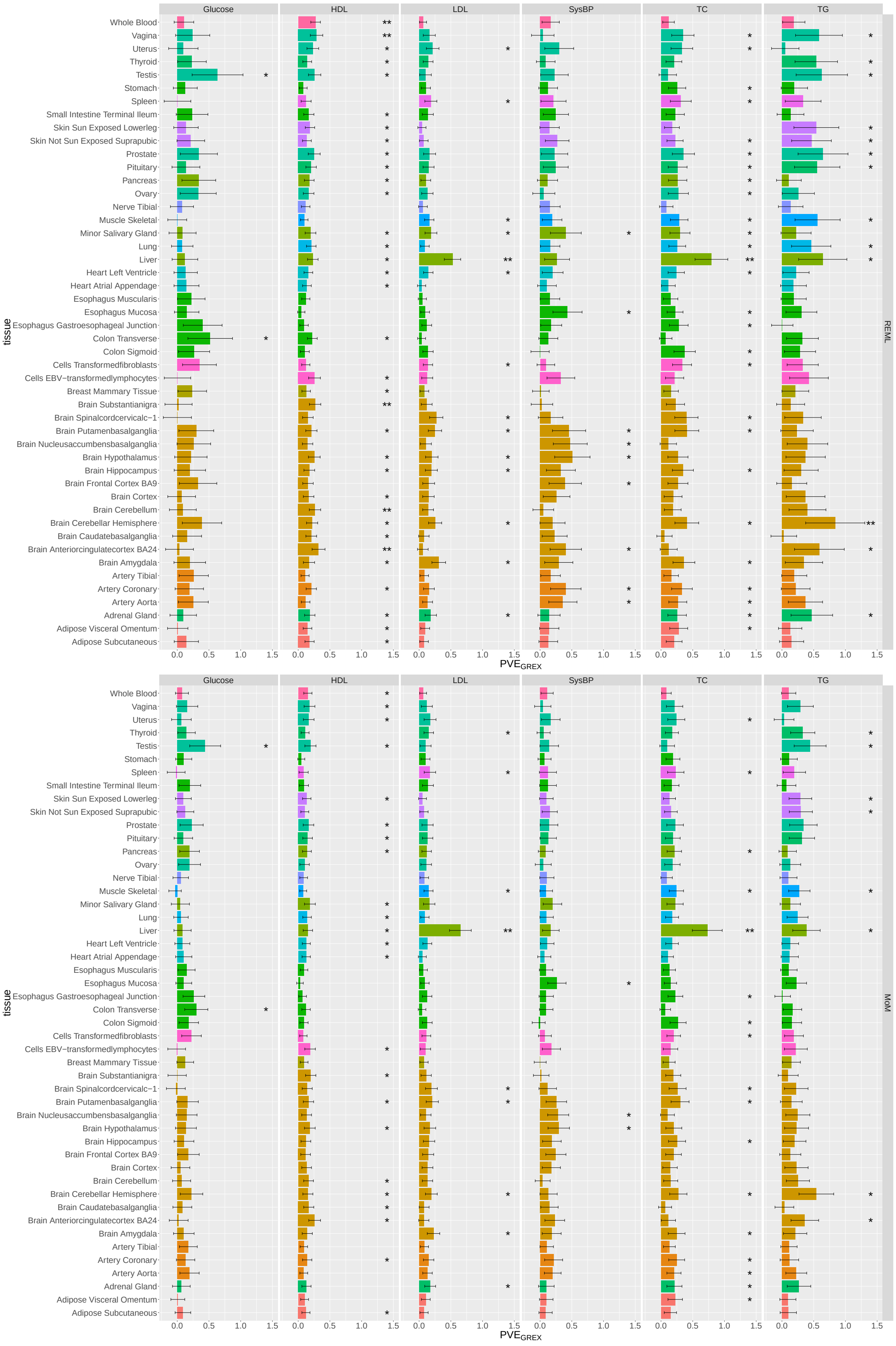

**Supplementary Figure 6:** REML and MoM estimates of $\text{PVE}_{\text{GREX}}/h_t^2$ for NFBC data.

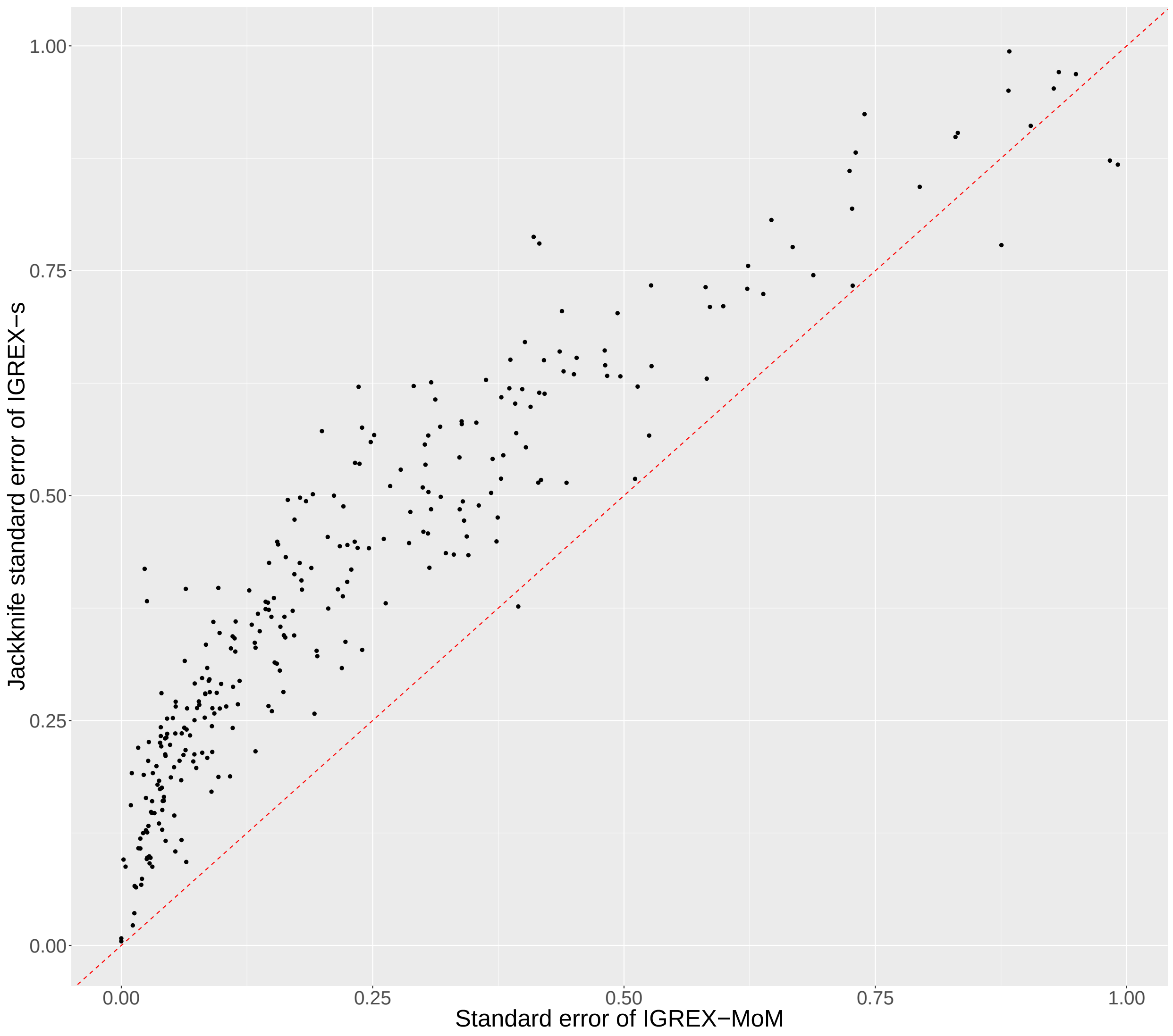

**Supplementary Figure 7:** Standard errors of $\text{PVE}_{\text{GREX}}$ for NFBC data estimated using block Jackknife and sandwich estimator.

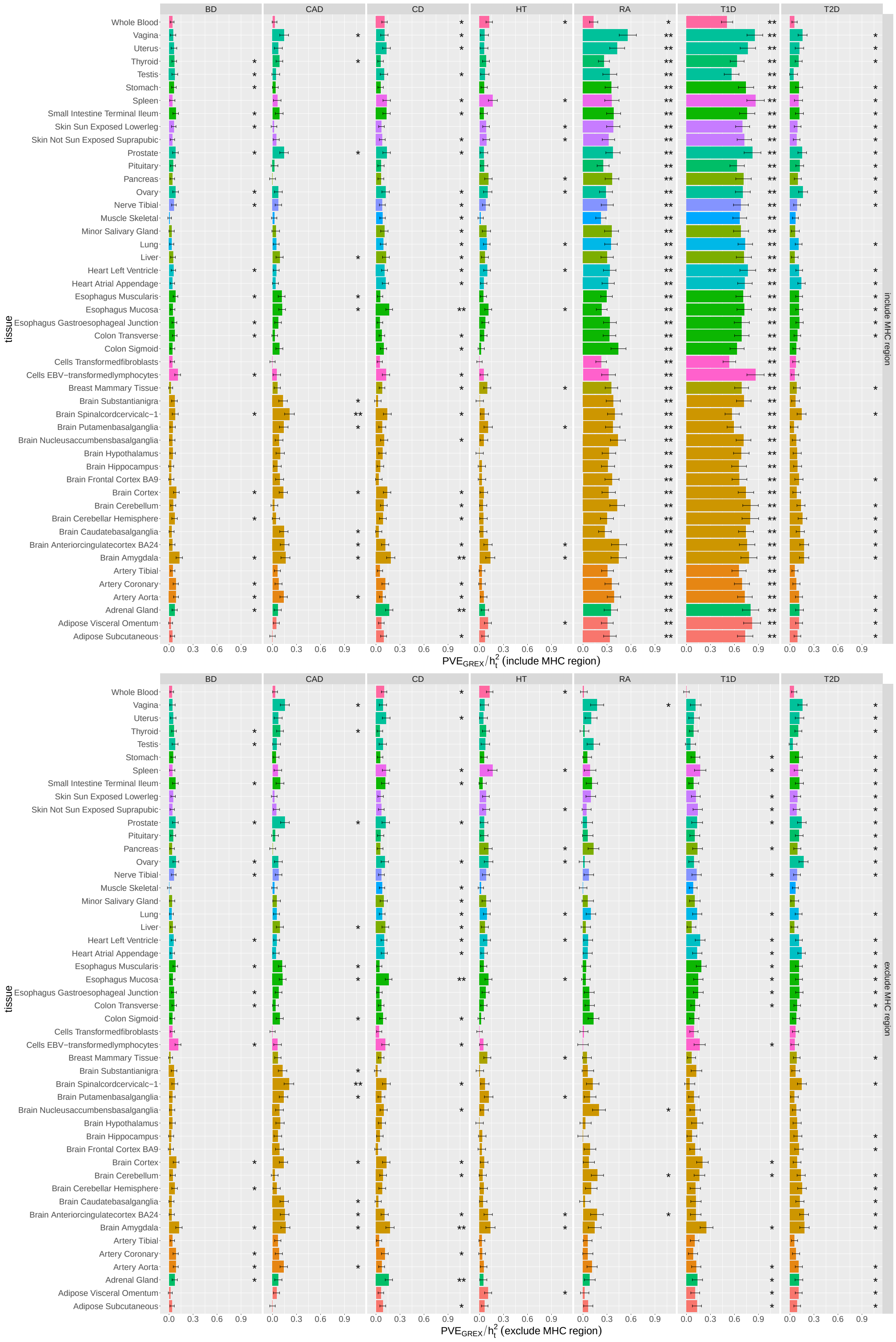

**Supplementary Figure 8:** REML estimates of $\mathrm{PVE}_{\mathrm{GREX}}/h_t^2$ for WTCCC data.

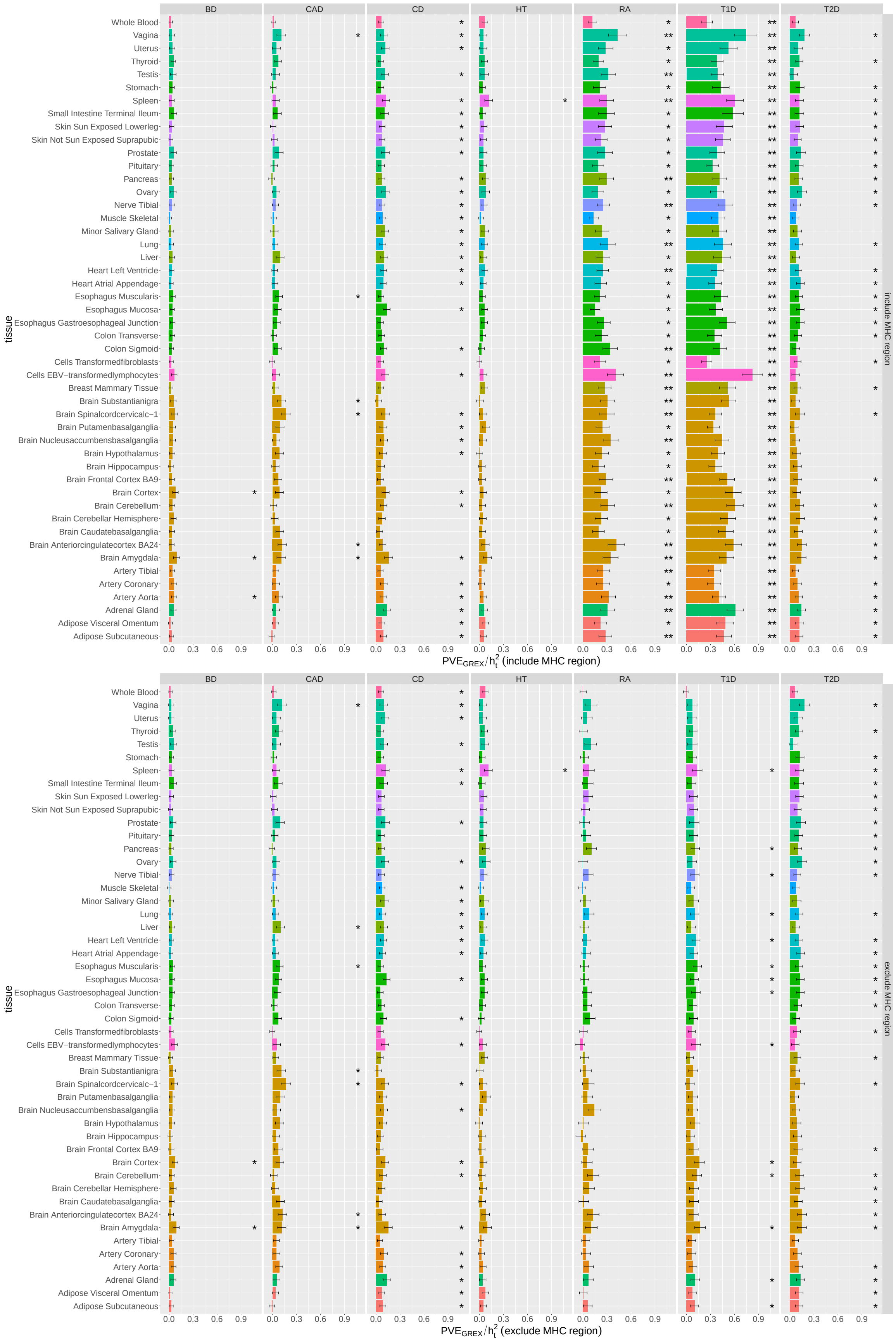

**Supplementary Figure 9:** MoM estimates of $\mathrm{PVE}_{\mathrm{GREX}}/h_t^2$ for WTCCC data.

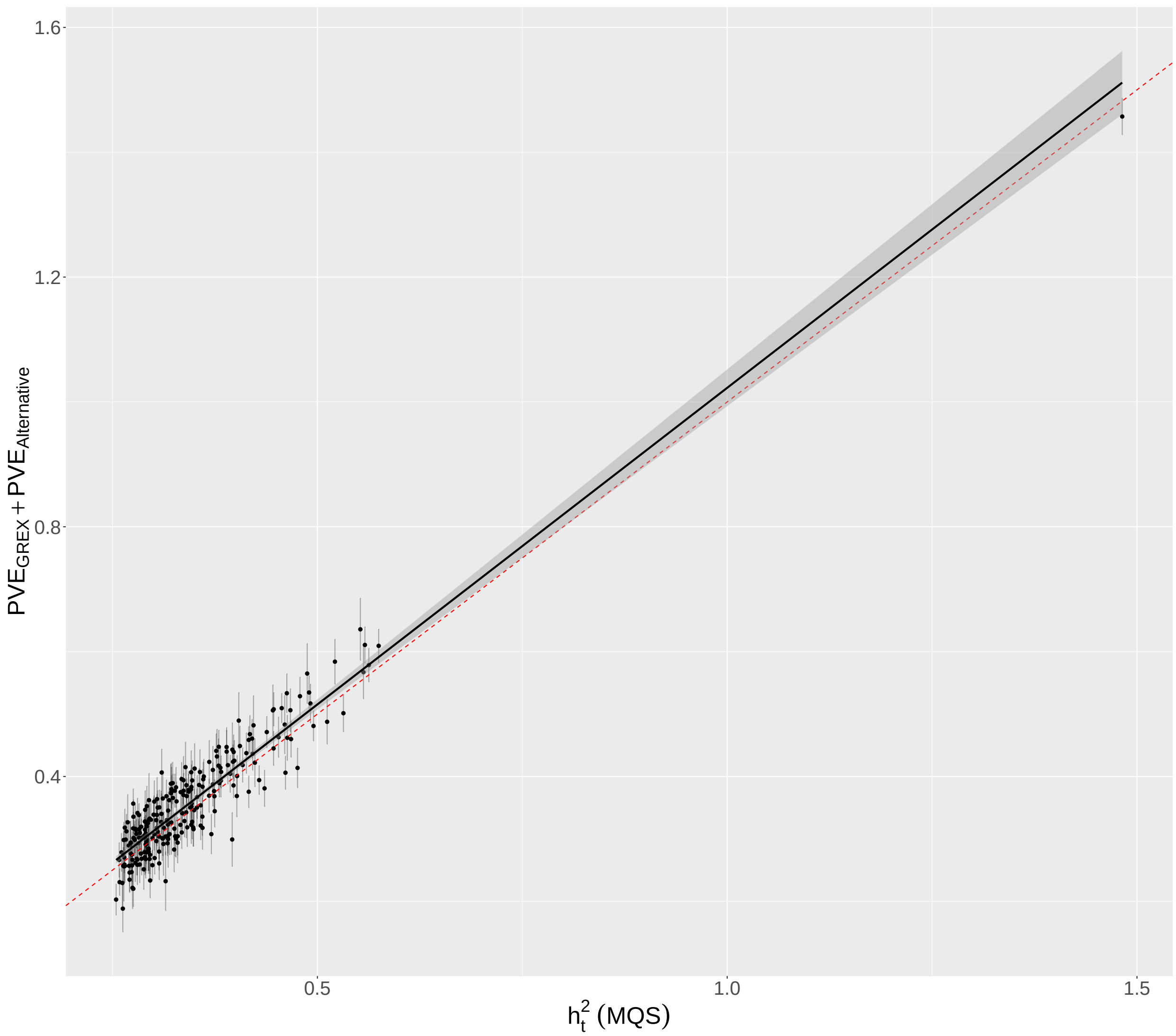

**Supplementary Figure 10:** Comparison of $\mathrm{PVE}_{\mathrm{GREX}} + \mathrm{PVE}_{\mathrm{Alternative}}$ estimated by IGREX and $h_t^2$ estimated by [1] in pQTL dataset.

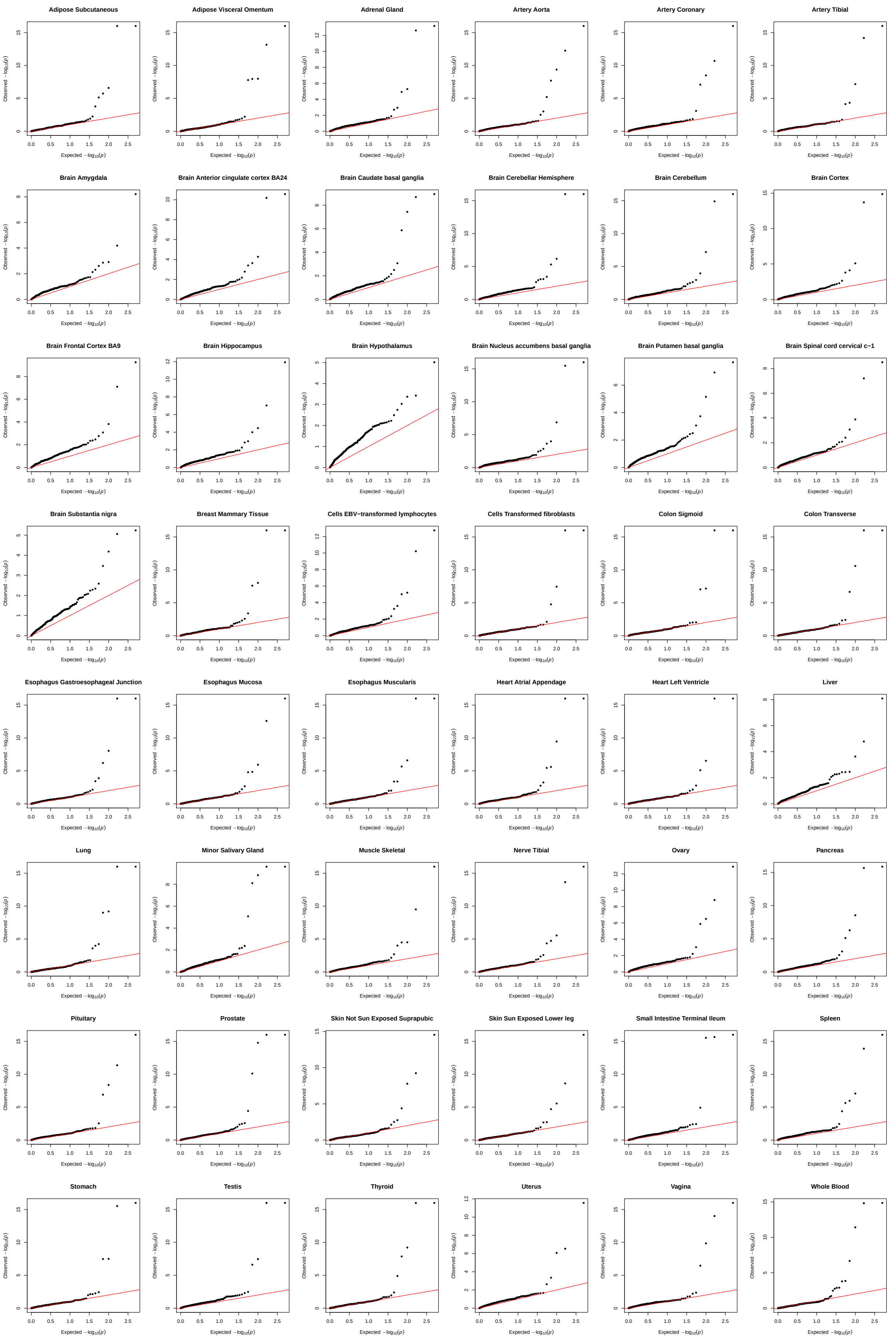

**Supplementary Figure 11:** QQ plot of $p$-values of $\text{PVE}_{\text{GREX}}$ for proteins in each of GTEx tissue.

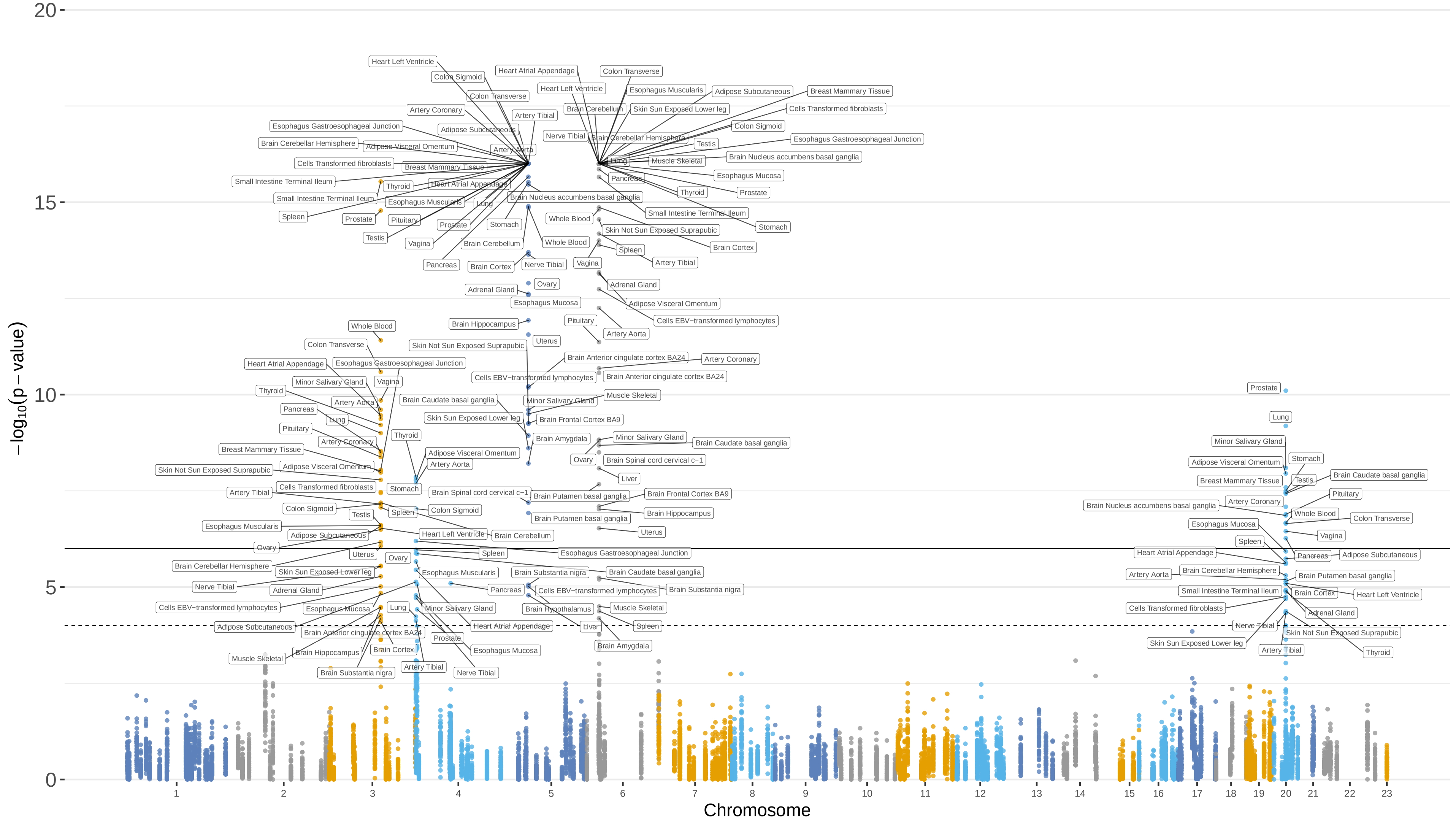

**Supplementary Figure 12:** Manhattan plot of protein-coding genes for pQTL study.

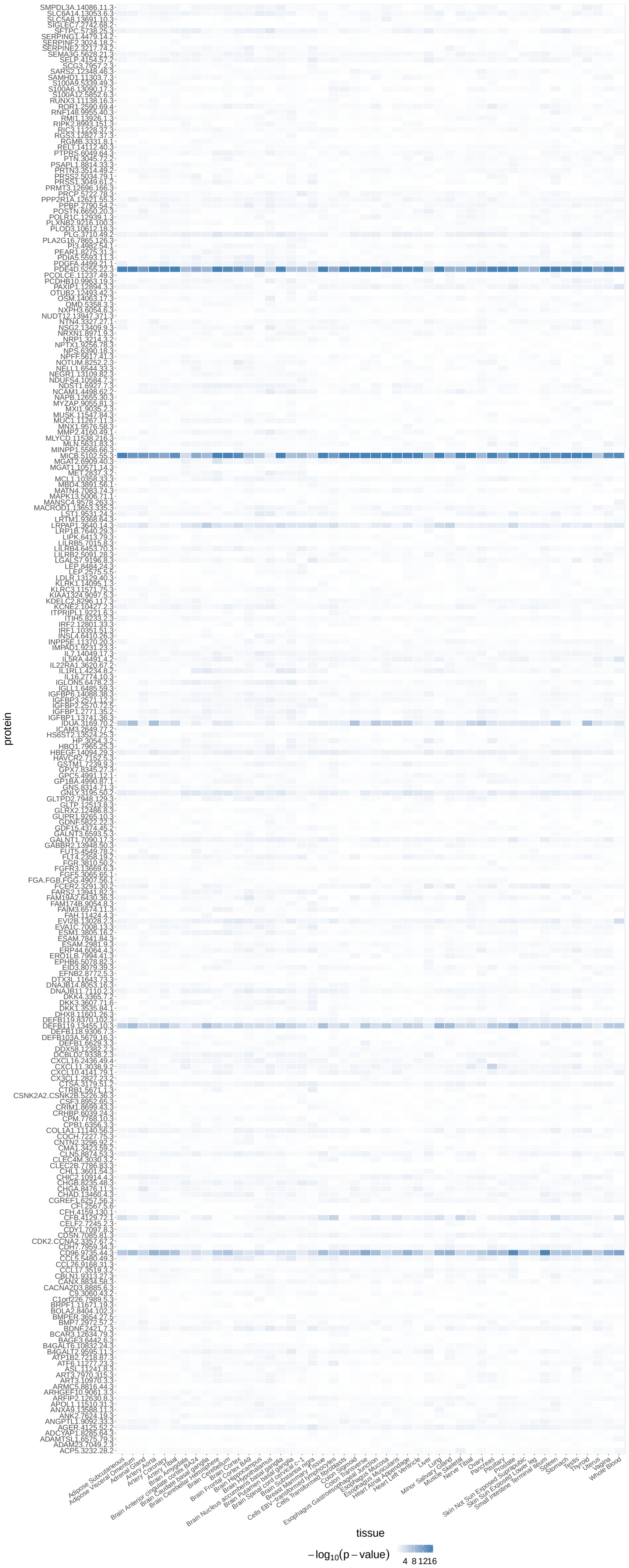

**Supplementary Figure 13:** Heatmap of $p$-values of $\text{PVE}_{\text{GREX}}$ for 249 proteins.

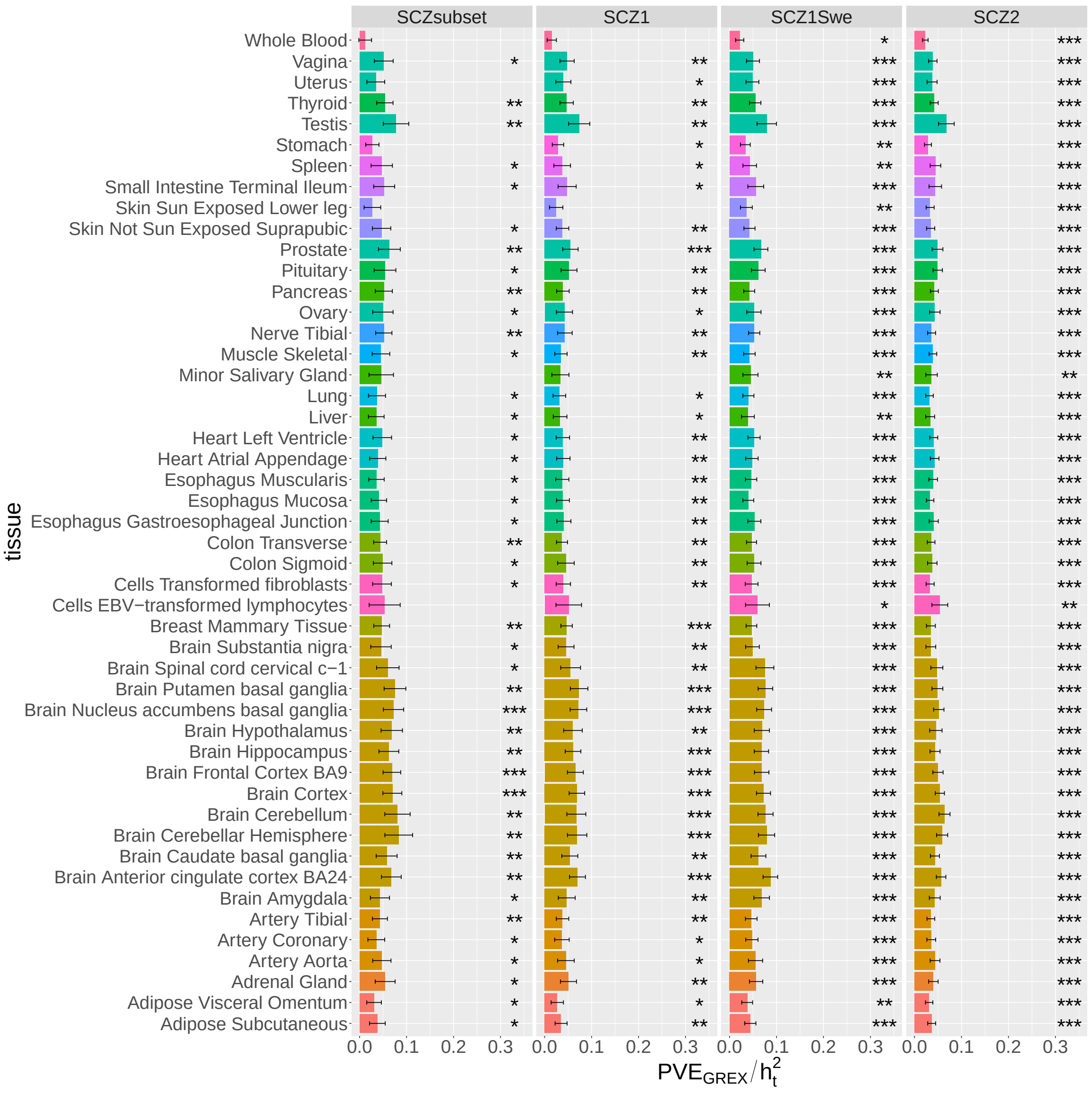

**Supplementary Figure 14:** Estimated $\mathrm{PVE}_{\mathrm{GREX}}/h_t^2$ for four independent schizophrenia datasets.

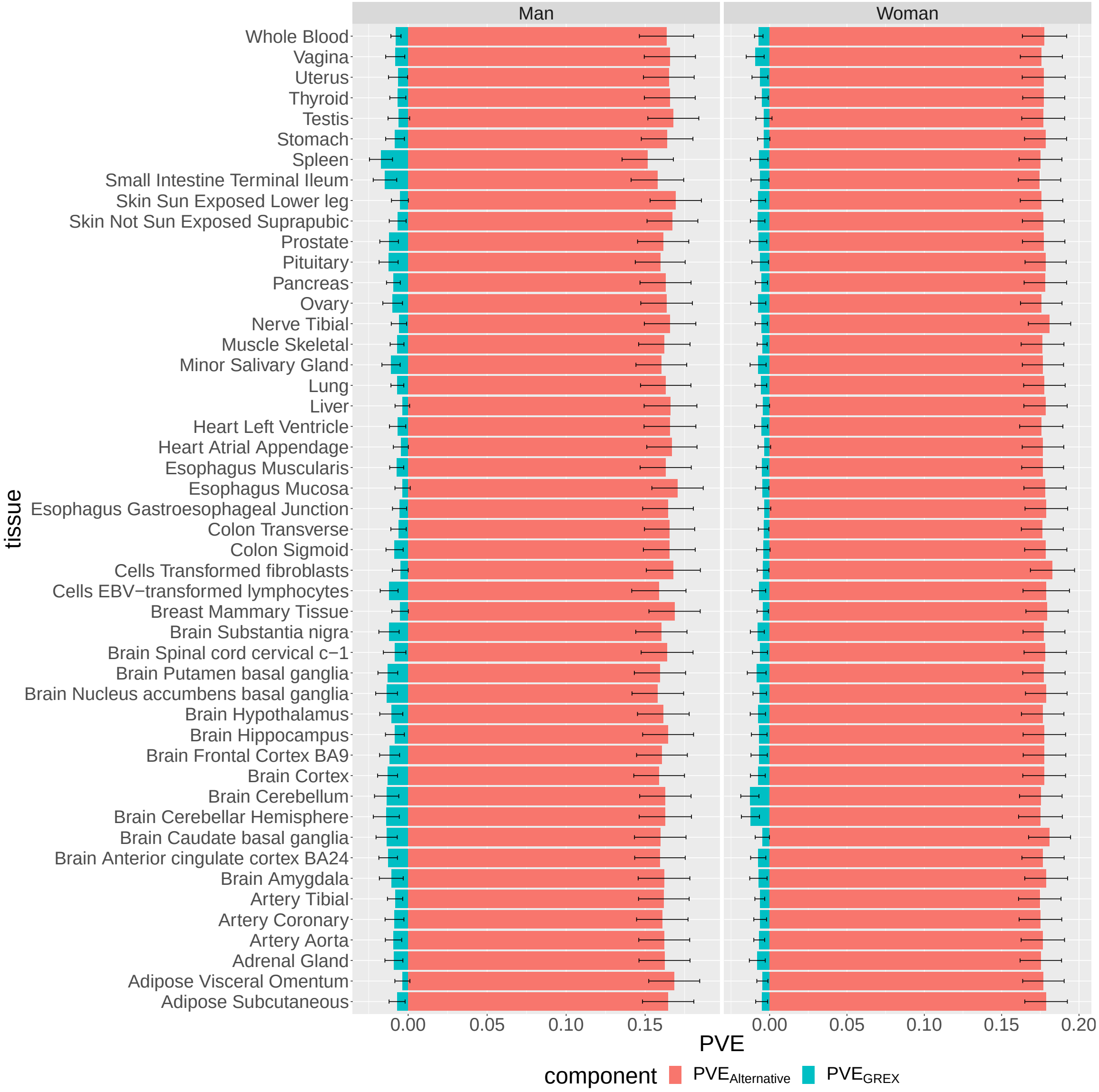

**Supplementary Figure 15:** Estimated $\mathrm{PVE}_{\mathrm{GREX}}$ and $\mathrm{PVE}_{\mathrm{Alternative}}$ for BMI datasets.

### 2 Supplementary Note

#### 2.1 IGREX model in the general form

IGREX can account for fixed effects in both eQTL and GWAS data. Let $\mathbf{W}_{r,g}$ be the set of $c_g$ covariates to be adjusted (e.g. age, sex, and the first few PCs) in the eQTL study of $g$-th gene and $\mathbf{W}$ be the set of $c$ covariates in the GWAS study. Then model (1) in the main text can be extended to incorporate the covariates:

$$\mathbf{y}_g = \mathbf{W}_{r,g}\mathbf{u}_g + \mathbf{X}_{r,g}\boldsymbol{\beta}_g + \mathbf{e}_{r,g}, \tag{S1}$$

where $\mathbf{u}_g \in \mathbb{R}^{c_g}$ is the fixed effect of $\mathbf{W}_{r,g} \in \mathbb{R}^{n_r \times c_g}$ and $\mathbf{e}_{r,g} \sim \mathcal{N}(0, \sigma^2_{r,g}\mathbf{I}_{n_r})$ is a vector of independent noise. Similarly, model (2) in main text becomes:

$$\mathbf{t} = \mathbf{W}\mathbf{v} + \sum_{g=1}^{G} \alpha_g \mathbf{X}_g \boldsymbol{\beta}_g + \mathbf{X}\boldsymbol{\gamma} + \boldsymbol{\epsilon}, \tag{S2}$$

where $\mathbf{v} \in \mathbb{R}^c$ is the fixed effect of $\mathbf{W} \in \mathbb{R}^{n \times c}$ and $\boldsymbol{\epsilon} \sim \mathcal{N}(0, \sigma^2_\epsilon \mathbf{I}_n)$ is a vector of independent noise. The prior distributions of effects $\boldsymbol{\beta}_g$, $\alpha_g$ and $\boldsymbol{\gamma}$ are given as:

$$\boldsymbol{\beta}_g \sim \mathcal{N}(\mathbf{0}, \sigma^2_{\beta_g}\mathbf{I}_{M_g}),\ \alpha_g \sim \mathcal{N}(\mathbf{0}, \sigma^2_\alpha),\ \boldsymbol{\gamma} \sim \mathcal{N}(\mathbf{0}, \sigma^2_\gamma \mathbf{I}_M).$$

The two-stage IGREX procedure first evaluates the posterior of $\boldsymbol{\beta}_g$ from the eQTL study based on model (S1) and then estimates $\text{PVE}_{\text{GREX}}$ from GWAS data by treating the obtained posterior as prior of $\boldsymbol{\beta}_g$ in model (S2).

#### 2.2 Stage-one: evaluate the posterior of $\boldsymbol{\beta}_g$ with PX-EM algorithm

At stage one, we evaluate the posterior $\boldsymbol{\beta}_g | \mathbf{y}_g, \mathbf{X}_{r,g}, \mathbf{W}_{r,g} \sim \mathcal{N}(\boldsymbol{\mu}_g, \boldsymbol{\Sigma}_g)$ from the eQTL data set $\mathcal{D}_{r,g} = \{\mathbf{y}_g, \mathbf{X}_{r,g}, \mathbf{W}_{r,g}\}$ corresponding to the $g$-th gene. This requires estimating the collection of parameters $\boldsymbol{\theta}_g = \{\mathbf{u}_g, \sigma^2_{\beta_g}, \sigma^2_{r,g}\}$ in model (S1). Here we derive an efficient PX-EM algorithm [2] to solve this linear mixed model. We first consider the parameter expanded version of (S1):

$$\mathbf{y}_g = \mathbf{W}_{r,g}\mathbf{u}_g + \delta \mathbf{X}_{r,g}\boldsymbol{\beta}_g + \mathbf{e}_{r,g},$$

where $\delta \in \mathbb{R}$ is the expanded parameter. The complete-data log-likelihood is given as

$$\begin{aligned} &\log \Pr(\mathbf{y}_g, \boldsymbol{\beta}_g | \boldsymbol{\theta}_g; \mathbf{W}_{r,g}, \mathbf{X}_{r,g}) \\ =& -\frac{n_r}{2}\log(2\pi\sigma^2_{r,g}) - \frac{1}{2\sigma^2_{r,g}}||\mathbf{y}_g - \mathbf{W}_{r,g}\mathbf{u}_g - \delta\mathbf{X}_{r,g}\boldsymbol{\beta}_g||^2 \\ &- \frac{M_g}{2}\log(2\pi\sigma^2_{\beta_g}) - \frac{1}{2\sigma^2_{\beta_g}}||\boldsymbol{\beta}_g||^2, \end{aligned} \tag{S3}$$

from which we can easily recognize that the the terms involving $\boldsymbol{\beta}_g$ are of a quadratic form:

$$\boldsymbol{\beta}_g^T \left( -\frac{\delta^2}{2\sigma^2_{r,g}}\mathbf{X}^T_{r,g}\mathbf{X}_{r,g} - \frac{1}{2\sigma^2_{\beta_g}}\mathbf{I}_{M_g} \right) \boldsymbol{\beta}_g + \frac{\delta}{\sigma^2_{r,g}}(\mathbf{y}_g - \mathbf{W}_{r,g}\mathbf{u}_g)^T \mathbf{X}_{r,g}\boldsymbol{\beta}_g + \text{Constant}.$$

Therefore, the posterior distribution of $\boldsymbol{\beta}_g$ is Gaussian $\mathcal{N}(\boldsymbol{\beta}_g|\boldsymbol{\mu}_g, \boldsymbol{\Sigma}_g)$, where

$$\begin{aligned} \boldsymbol{\Sigma}_g^{-1} &= \frac{\delta^2}{\sigma^2_{r,g}}\mathbf{X}^T_{r,g}\mathbf{X}_{r,g} + \frac{1}{\sigma^2_{\beta_g}}\mathbf{I}_{M_g}, \\ \boldsymbol{\mu}_g &= \left( \frac{\delta^2}{\sigma^2_{r,g}}\mathbf{X}^T_{r,g}\mathbf{X}_{r,g} + \frac{1}{\sigma^2_{\beta_g}}\mathbf{I}_{M_g} \right)^{-1} \frac{\delta}{\sigma^2_{r,g}}\mathbf{X}^T_{r,g}(\mathbf{y}_g - \mathbf{W}_{r,g}\mathbf{u}_g). \end{aligned}$$

Now in the E-step, we evaluate the $\mathcal{Q}$-function by taking the expectation of the complete-data log-likelihood (S3) with respect to the posterior $\mathcal{N}(\boldsymbol{\beta}_g|\boldsymbol{\mu}_g, \boldsymbol{\Sigma}_g)$. Specifically, the quadratic terms involving $\boldsymbol{\beta}_g$ in (S3) are evaluated as following:

$$\begin{aligned}\mathbb{E}[||\tilde{\mathbf{y}}_g - \delta\mathbf{X}_{r,g}\boldsymbol{\beta}_g||^2] &= \mathbb{E}[\tilde{\mathbf{y}}_g^T\tilde{\mathbf{y}}_g - 2\delta\tilde{\mathbf{y}}_g^T\mathbf{X}_{r,g}\boldsymbol{\beta}_g + \delta^2\boldsymbol{\beta}_g^T\mathbf{X}_{r,g}^T\mathbf{X}_{r,g}\boldsymbol{\beta}_g] \\ &= \tilde{\mathbf{y}}_g^T\tilde{\mathbf{y}}_g - 2\delta\tilde{\mathbf{y}}_g^T\mathbf{X}_{r,g}\boldsymbol{\mu}_g + \delta^2\mathrm{tr}(\mathbf{X}_{r,g}^T\mathbf{X}_{r,g}\boldsymbol{\Sigma}_g), \\ \mathbb{E}[||\boldsymbol{\beta}_g||^2] &= \boldsymbol{\mu}_g^T\boldsymbol{\mu}_g + \mathrm{tr}(\boldsymbol{\Sigma}_g),\end{aligned}$$

where $\tilde{\mathbf{y}}_g = \mathbf{y}_g - \mathbf{W}_{r,g}\mathbf{u}_g$. Then the $\mathcal{Q}$-function given the current parameter estimates $\boldsymbol{\theta}_{g,old}$ is obtained as:

$$\begin{aligned}\mathcal{Q}(\boldsymbol{\theta}_g|\boldsymbol{\theta}_{g,old}) = &-\frac{n_r}{2}\log(2\pi\sigma_{r,g}^2) - \frac{M_g}{2}\log(2\pi\sigma_{\beta_g}^2) \\ &- \frac{1}{2\sigma_{r,g}^2}||\mathbf{y}_g - \mathbf{W}_{r,g}\mathbf{u}_g - \delta\mathbf{X}_{r,g}\boldsymbol{\mu}_g||^2 - \frac{1}{2\sigma_{\beta_g}^2}||\boldsymbol{\mu}_g||^2 \\ &- \frac{1}{2\sigma_{\beta_g}^2}\boldsymbol{\mu}_g^T\boldsymbol{\mu}_g - \mathrm{tr}\left(\left(\frac{\delta^2}{2\sigma_{r,g}^2}\mathbf{X}_{r,g}^T\mathbf{X}_{r,g} + \frac{1}{2\sigma_{\beta_g}^2}\mathbf{I}_{M_g}\right)\boldsymbol{\Sigma}_g\right).\end{aligned}$$

It the M-step, the new estimates of parameter $\boldsymbol{\theta}_g$ is obtained by setting the derivative of $\mathcal{Q}$-function to be zero. The resulting updates are given as follows:

$$\begin{aligned}\delta &= \frac{(\mathbf{y}_g - \mathbf{W}_{r,g}\mathbf{u}_g)^T\mathbf{X}_{r,g}\boldsymbol{\mu}_g}{\boldsymbol{\mu}_g^T\mathbf{X}_{r,g}^T\mathbf{X}_{r,g}\boldsymbol{\mu}_g + \mathrm{tr}(\mathbf{X}_{r,g}^T\mathbf{X}_{r,g}\boldsymbol{\Sigma}_g)}, \\ \mathbf{u}_g &= (\mathbf{X}_{r,g}^T\mathbf{X}_{r,g})^{-1}\mathbf{X}_{r,g}^T(\mathbf{y}_g - \delta\mathbf{X}_{r,g}\boldsymbol{\mu}_g), \\ \sigma_{r,g}^2 &= \frac{1}{n_r}[||\mathbf{y}_g - \mathbf{W}_{r,g}\mathbf{u}_g - \delta\mathbf{X}_{r,g}\boldsymbol{\mu}_g||^2 + \delta^2\mathrm{tr}(\mathbf{X}_{r,g}^T\mathbf{X}_{r,g}\boldsymbol{\Sigma}_g)], \\ \sigma_{\beta_g}^2 &= \frac{1}{M_g}[\boldsymbol{\mu}_g^T\boldsymbol{\mu}_g + \mathrm{tr}(\boldsymbol{\Sigma}_g)].\end{aligned}$$

This PX-EM algorithm is summarized in Algorithm 1. After convergence, the posterior mean and variance of $\mathcal{N}(\boldsymbol{\beta}_g|\boldsymbol{\mu}_g, \boldsymbol{\Sigma}_g)$ can be evaluated given the obtained parameter estimates $\hat{\boldsymbol{\theta}}_g = \{\hat{\mathbf{u}}_g, \hat{\sigma}_{\beta_g}^2, \hat{\sigma}_{r,g}^2\}$:

$$\begin{aligned}\boldsymbol{\Sigma}_g^{-1} &= \frac{1}{\hat{\sigma}_{r,g}^2}\mathbf{X}_{r,g}^T\mathbf{X}_{r,g} + \frac{1}{\hat{\sigma}_{\beta_g}^2}\mathbf{I}_{M_g}, \\ \boldsymbol{\mu}_g &= \left(\frac{1}{\hat{\sigma}_{r,g}^2}\mathbf{X}_{r,g}^T\mathbf{X}_{r,g} + \frac{1}{\hat{\sigma}_{\beta_g}^2}\mathbf{I}_{M_g}\right)^{-1}\frac{1}{\hat{\sigma}_{r,g}^2}\mathbf{X}_{r,g}^T(\mathbf{y}_g - \mathbf{W}_{r,g}\hat{\mathbf{u}}_g).\end{aligned} \tag{S4}$$

#### 2.3 Stage-two: estimate $\mathrm{PVE}_{\mathrm{GREX}}$

Given the parameters in (S4), we then estimate $\mathrm{PVE}_{\mathrm{GREX}} = \frac{\mathrm{Var}(\sum_{g=1}^{G}\alpha_g\mathbf{x}_g^T\boldsymbol{\beta}_g)}{\mathrm{Var}(t)}$ by treating the posterior distribution $\mathcal{N}(\boldsymbol{\beta}_g|\boldsymbol{\mu}_g, \boldsymbol{\Sigma}_g)$ as prior distribution of $\boldsymbol{\beta}_g$ in model (S2). To account

**Algorithm 1** PX-EM algorithm for model (S1)

Initialization: Parameters are initialized by setting $\mathbf{u}_g = (\mathbf{W}_{r,g}^T\mathbf{W}_{r,g})^{-1}\mathbf{W}_{r,g}^T\mathbf{y}_g$, $\sigma_{r,g}^2 = \sigma_{\beta_g}^2 = \mathrm{Var}(y - \mathbf{W}_{r,g}\mathbf{u}_g)/2$.

**repeat**

**E-step:** At the $t$-th iteration, evaluate the posterior $\mathcal{N}(\boldsymbol{\beta}_g|\boldsymbol{\mu}_g, \boldsymbol{\Sigma}_g)$ given the current parameter estimates $\boldsymbol{\theta}_g^{(t)} = \{\mathbf{u}_g^{(t)}, (\sigma_{r,g}^{(t)})^2, (\sigma_{\beta_g}^{(t)})^2\}$ and $\delta^{(t)} = 1$:

$$\boldsymbol{\Sigma}_g^{-1} = \frac{(\delta^{(t)})^2}{(\sigma_{r,g}^{(t)})^2}\mathbf{X}_{r,g}^T\mathbf{X}_{r,g} + \frac{1}{(\sigma_{\beta_g}^{(t)})^2}\mathbf{I}_{M_g},$$

$$\boldsymbol{\mu}_g = \left(\frac{(\delta^{(t)})^2}{(\sigma_{r,g}^{(t)})^2}\mathbf{X}_{r,g}^T\mathbf{X}_{r,g} + \frac{1}{(\sigma_{\beta_g}^{(t)})^2}\mathbf{I}_{M_g}\right)^{-1}\frac{(\delta^{(t)})^2}{(\sigma_{r,g}^{(t)})^2}\mathbf{X}_{r,g}^T(\mathbf{y}_g - \mathbf{W}_{r,g}\mathbf{u}_g^{(t)})$$

**M-step:** Update the model parameters $\boldsymbol{\theta}_g$ by

$$\delta^{(t+1)} = \frac{(\mathbf{y}_g - \mathbf{W}_{r,g}\mathbf{u}_g^{(t)})^T\mathbf{X}_{r,g}\boldsymbol{\mu}_g}{\boldsymbol{\mu}_g^T\mathbf{X}_{r,g}^T\mathbf{X}_{r,g}\boldsymbol{\mu}_g + \mathrm{tr}(\mathbf{X}_{r,g}^T\mathbf{X}_{r,g}\boldsymbol{\Sigma}_g)},$$

$$\mathbf{u}_g^{(t+1)} = (\mathbf{X}_{r,g}^T\mathbf{X}_{r,g})^{-1}\mathbf{X}_{r,g}^T(\mathbf{y}_g - \delta\mathbf{X}_{r,g}\boldsymbol{\mu}_g),$$

$$(\sigma_{r,g}^{(t+1)})^2 = \frac{1}{n_r}[||\mathbf{y}_g - \mathbf{W}_{r,g}\mathbf{u}_g^{(t)} - \delta\mathbf{X}_{r,g}\boldsymbol{\mu}_g||^2 + \delta^2\mathrm{tr}(\mathbf{X}_{r,g}^T\mathbf{X}_{r,g}\boldsymbol{\Sigma}_g)],$$

$$(\sigma_{\beta_g}^{(t+1)})^2 = \frac{1}{M_g}[\boldsymbol{\mu}_g^T\boldsymbol{\mu}_g + \mathrm{tr}(\boldsymbol{\Sigma}_g)].$$

**Reduction-step:** Rescale $(\sigma_{\beta_g}^{(t+1)})^2 = (\delta^{(t+1)})^2(\sigma_{\beta_g}^{(t+1)})^2$ and reset $\delta^{(t+1)} = 1$.

**until** the incomplete-data log-likelihood stops increasing or maximum iteration is reached for the fixed effects $\mathbf{W}$, we first multiply the projection matrix $\mathbf{M} = \mathbf{I}_n - \mathbf{W}(\mathbf{W}^T\mathbf{W})^{-1}\mathbf{W}^T$ on both sides of model (S2). Then, we have $\mathbb{E}(\mathbf{Mt}|\boldsymbol{\alpha}) = \sum_{g=1}^{G}\alpha_g\mathbf{MX}_g\boldsymbol{\mu}_g$ and $\mathrm{Cov}(\mathbf{t}|\boldsymbol{\alpha}) = \sum_{g=1}^{G}\alpha_g^2\mathbf{MX}_g\boldsymbol{\Sigma}_g(\mathbf{MX}_g)^T + \sigma_\gamma^2\mathbf{MX}(\mathbf{MX})^T + \sigma_\epsilon^2\mathbf{M}$. By applying the law of total expectation and total variance, we obtain $\mathbb{E}(\mathbf{Mt}) = \mathbb{E}(\mathbb{E}(\mathbf{Mt}|\boldsymbol{\alpha})) = \mathbf{0}$ and

$$\mathrm{Cov}(\mathbf{Mt}) = \mathrm{Cov}(\mathbb{E}(\mathbf{Mt}|\boldsymbol{\alpha})) + \mathbb{E}(\mathrm{Cov}(\mathbf{Mt}|\boldsymbol{\alpha})) = \sigma_\alpha^2\mathbf{MK}_\alpha\mathbf{M} + \sigma_\gamma^2\mathbf{MK}_\gamma\mathbf{M} + \sigma_\epsilon^2\mathbf{M} =: \boldsymbol{\Omega},$$

respectively, where $\mathbf{K}_\alpha = \sum_{g=1}^{G}\mathbf{X}_g(\boldsymbol{\mu}_g\boldsymbol{\mu}_g^T + \boldsymbol{\Sigma}_g)\mathbf{X}_g^T$ and $\mathbf{K}_\gamma = \mathbf{XX}^T$. Clearly, the $i$-th diagonal element of $\sigma_\alpha^2\mathbf{MK}_\alpha\mathbf{M}$ and $\sigma_\gamma^2\mathbf{MK}_\gamma\mathbf{M}$ represents the variance explained by GREX and alternative genetic effects, respectively. Hence, the $\mathrm{PVE}_{\mathrm{GREX}}$, $\mathrm{PVE}_{\mathrm{Alternative}}$, $h_t^2$ and $\mathrm{PVE}_{\mathrm{GREX}}/h_t^2$ are estimated by

$$\begin{aligned}
\widehat{\mathrm{PVE}}_{\mathrm{GREX}} &= \frac{\mathrm{tr}(\hat{\sigma}_\alpha^2\mathbf{MK}_\alpha\mathbf{M})}{\mathrm{tr}(\hat{\sigma}_\alpha^2\mathbf{MK}_\alpha\mathbf{M} + \hat{\sigma}_\gamma^2\mathbf{MK}_\gamma\mathbf{M} + \hat{\sigma}_\epsilon^2\mathbf{M})},\\
\widehat{\mathrm{PVE}}_{\mathrm{Alternative}} &= \frac{\mathrm{tr}(\hat{\sigma}_\gamma^2\mathbf{MK}_\gamma\mathbf{M})}{\mathrm{tr}(\hat{\sigma}_\alpha^2\mathbf{MK}_\alpha\mathbf{M} + \hat{\sigma}_\gamma^2\mathbf{MK}_\gamma\mathbf{M} + \hat{\sigma}_\epsilon^2\mathbf{M})},\\
\hat{h}_t^2 &= \frac{\mathrm{tr}(\hat{\sigma}_\alpha^2\mathbf{MK}_\alpha\mathbf{M} + \hat{\sigma}_\gamma^2\mathbf{MK}_\gamma\mathbf{M})}{\mathrm{tr}(\hat{\sigma}_\alpha^2\mathbf{MK}_\alpha\mathbf{M} + \hat{\sigma}_\gamma^2\mathbf{MK}_\gamma\mathbf{M} + \hat{\sigma}_\epsilon^2\mathbf{M})},\\
\widehat{\mathrm{PVE}}_{\mathrm{GREX}}/\hat{h}_t^2 &= \frac{\mathrm{tr}(\hat{\sigma}_\alpha^2\mathbf{MK}_\alpha\mathbf{M})}{\mathrm{tr}(\hat{\sigma}_\alpha^2\mathbf{MK}_\alpha\mathbf{M} + \hat{\sigma}_\gamma^2\mathbf{MK}_\gamma\mathbf{M})},
\end{aligned} \tag{S5}$$

where $\hat{\sigma}_\alpha^2$, $\hat{\sigma}_\gamma^2$ and $\hat{\sigma}_\epsilon^2$ are the estimated values of $\sigma_\alpha^2$, $\sigma_\gamma^2$ and $\sigma_\epsilon^2$, respectively. Letting $\boldsymbol{\theta} = \{\sigma_\alpha^2, \sigma_\gamma^2, \sigma_\epsilon^2\}$ be the collection of parameters to be estimated, its corresponding estimate $\hat{\boldsymbol{\theta}} = \{\hat{\sigma}_\alpha^2, \hat{\sigma}_\gamma^2, \hat{\sigma}_\epsilon^2\}$ can be obtained by REML or MoM when the individual-level GWAS data $\mathcal{D}_i = \{\mathbf{t}, \mathbf{X}, \mathbf{W}\}$ is available. Both approaches are provided in the IGREX framework (IGREX-i): REML is statistically more efficient by assuming normality of $\mathbf{t}$ while MoM is more robust and computationally efficient. Besides, we developed IGREX-s based on MoM to handle GWAS summary statistics when the individual-level GWAS data is inaccessible.

##### 2.3.1 IGREX-i

The MoM estimate of $\boldsymbol{\psi}$ is obtained by minimizing the distance between the second moment of $\mathbf{t}$ at the population level and that at the sample level:

$$\min_{\boldsymbol{\psi}} \; f(\boldsymbol{\psi}) = ||(\mathbf{Mt})(\mathbf{Mt})^T - (\sigma_\alpha^2\mathbf{MK}_\alpha\mathbf{M} + \sigma_\gamma^2\mathbf{MK}_\gamma\mathbf{M} + \sigma_\epsilon^2\mathbf{M})||^2.$$

The solution of this optimization problem is obtained by setting $\frac{\partial f(\boldsymbol{\psi})}{\partial\sigma_\alpha^2} = \frac{\partial f(\boldsymbol{\psi})}{\partial\sigma_\gamma^2} = \frac{\partial f(\boldsymbol{\psi})}{\partial\sigma_\epsilon^2} = 0$, which gives the estimating equation:

$$\mathbf{S}\boldsymbol{\psi} = \mathbf{q}, \tag{S6}$$

$$\text{with } \mathbf{S} = \begin{bmatrix} \text{tr}((\mathbf{MK}_\alpha)^2) & \text{tr}(\mathbf{MK}_\alpha\mathbf{MK}_\gamma) & \text{tr}(\mathbf{MK}_\alpha) \\ \text{tr}(\mathbf{MK}_\alpha\mathbf{MK}_\gamma) & \text{tr}((\mathbf{MK}_\gamma)^2) & \text{tr}(\mathbf{MK}_\gamma) \\ \text{tr}(\mathbf{MK}_\alpha) & \text{tr}(\mathbf{MK}_\gamma) & n-c \end{bmatrix},\ \boldsymbol{\psi} = \begin{bmatrix} \sigma_\alpha^2 \\ \sigma_\gamma^2 \\ \sigma_\epsilon^2 \end{bmatrix},\ \mathbf{q} = \begin{bmatrix} \mathbf{t}^T\mathbf{MK}_\alpha\mathbf{Mt} \\ \mathbf{t}^T\mathbf{MK}_\gamma\mathbf{Mt} \\ \mathbf{t}^T\mathbf{Mt} \end{bmatrix}.$$

Thus, $\hat{\boldsymbol{\psi}}$ is obtained by calculating $\hat{\boldsymbol{\psi}} = \mathbf{S}^{-1}\mathbf{q}$. To estimate the standard error of $\widehat{\text{PVE}}_{\text{GREX}}$, we first calculate the standard errors of $\hat{\boldsymbol{\psi}}$ using sandwich estimator:

$$\text{Cov}(\hat{\boldsymbol{\psi}}) = \mathbf{S}^{-1}\text{Cov}(\mathbf{q})\mathbf{S}^{-1},$$

where

$$\begin{aligned} \text{Cov}(\mathbf{q}) &= \begin{bmatrix} \text{Var}(\mathbf{t}^T\mathbf{MK}_\alpha\mathbf{Mt}) & . & . \\ \text{Cov}(\mathbf{t}^T\mathbf{MK}_\alpha\mathbf{Mt}, \mathbf{t}^T\mathbf{MK}_\gamma\mathbf{Mt}) & \text{Var}(\mathbf{t}^T\mathbf{MK}_\gamma\mathbf{Mt}) & . \\ \text{Cov}(\mathbf{t}^T\mathbf{MK}_\alpha\mathbf{Mt}, \mathbf{t}^T\mathbf{t}) & \text{Cov}(\mathbf{t}^T\mathbf{MK}_\gamma\mathbf{Mt}, \mathbf{t}^T\mathbf{Mt}) & \text{Var}(\mathbf{t}^T\mathbf{Mt}) \end{bmatrix} \\ &= \begin{bmatrix} 2\text{tr}((\mathbf{MK}_\alpha\mathbf{M\Omega})^2) & . & . \\ 2\text{tr}(\mathbf{MK}_\alpha\mathbf{MK}_\gamma(\mathbf{M\Omega})^2) & 2\text{tr}((\mathbf{MK}_\gamma\mathbf{M\Omega})^2) & . \\ 2\text{tr}(\mathbf{MK}_\alpha(\mathbf{M\Omega})^2) & 2\text{tr}(\mathbf{MK}_\gamma(\mathbf{M\Omega})^2) & 2\text{tr}((\mathbf{M\Omega})^2) \end{bmatrix} \end{aligned}$$

is the symmetric covariance matrix of $\mathbf{q}$. Then, the variances of the PVE estimators defined in (S5) are obtained by the delta method:

$$\begin{aligned} \text{Var}(\widehat{\text{PVE}}_{\text{GREX}}) &= \nabla_G^T\text{Cov}(\hat{\boldsymbol{\psi}})\nabla_G \\ \text{Var}(\widehat{\text{PVE}}_{\text{Alternative}}) &= \nabla_A^T\text{Cov}(\hat{\boldsymbol{\psi}})\nabla_A \\ \text{Var}(\hat{h}_t^2) &= \nabla_h^T\text{Cov}(\hat{\boldsymbol{\psi}})\nabla_h \\ \text{Var}(\widehat{\text{PVE}}_{\text{GREX}}/\hat{h}_t^2) &= \nabla_{prop}^T\text{Cov}(\hat{\boldsymbol{\psi}})\nabla_{prop}, \end{aligned}$$

where

$$\nabla_G = \begin{bmatrix} \frac{\hat{\sigma}_\gamma^2\text{tr}(\mathbf{MK}_\alpha)\text{tr}(\mathbf{MK}_\gamma)+(n-c)\hat{\sigma}_\epsilon^2\text{tr}(\mathbf{MK}_\alpha)}{(\hat{\sigma}_\alpha^2\text{tr}(\mathbf{MK}_\alpha)+\hat{\sigma}_\gamma^2\text{tr}(\mathbf{MK}_\gamma)+\hat{\sigma}_\epsilon^2(n-c))^2} \\ -\frac{\hat{\sigma}_\alpha^2\text{tr}(\mathbf{MK}_\alpha)\text{tr}(\mathbf{MK}_\gamma)}{(\hat{\sigma}_\alpha^2\text{tr}(\mathbf{MK}_\alpha)+\hat{\sigma}_\gamma^2\text{tr}(\mathbf{MK}_\gamma)+\hat{\sigma}_\epsilon^2(n-c))^2} \\ -\frac{(n-c)\hat{\sigma}_\alpha^2\text{tr}(\mathbf{MK}_\alpha)}{(\hat{\sigma}_\alpha^2\text{tr}(\mathbf{MK}_\alpha)+\hat{\sigma}_\gamma^2\text{tr}(\mathbf{MK}_\gamma)+\hat{\sigma}_\epsilon^2(n-c))^2} \end{bmatrix},$$

$$\nabla_A = \begin{bmatrix} -\frac{\hat{\sigma}_\gamma^2\text{tr}(\mathbf{MK}_\alpha)\text{tr}(\mathbf{MK}_\gamma)}{(\hat{\sigma}_\alpha^2\text{tr}(\mathbf{MK}_\alpha)+\hat{\sigma}_\gamma^2\text{tr}(\mathbf{MK}_\gamma)+\hat{\sigma}_\epsilon^2(n-c))^2} \\ \frac{\hat{\sigma}_\alpha^2\text{tr}(\mathbf{MK}_\alpha)\text{tr}(\mathbf{MK}_\gamma)+(n-c)\hat{\sigma}_\epsilon^2\text{tr}(\mathbf{MK}_\gamma)}{(\hat{\sigma}_\alpha^2\text{tr}(\mathbf{MK}_\alpha)+\hat{\sigma}_\gamma^2\text{tr}(\mathbf{MK}_\gamma)+\hat{\sigma}_\epsilon^2(n-c))^2} \\ -\frac{(n-c)\hat{\sigma}_\gamma^2\text{tr}(\mathbf{MK}_\gamma)}{(\hat{\sigma}_\alpha^2\text{tr}(\mathbf{MK}_\alpha)+\hat{\sigma}_\gamma^2\text{tr}(\mathbf{MK}_\gamma)+\hat{\sigma}_\epsilon^2(n-c))^2} \end{bmatrix},$$

$$\nabla_h = \begin{bmatrix} \frac{(n-c)\hat{\sigma}_\epsilon^2\text{tr}(\mathbf{MK}_\alpha)}{(\hat{\sigma}_\alpha^2\text{tr}(\mathbf{MK}_\alpha)+\hat{\sigma}_\gamma^2\text{tr}(\mathbf{MK}_\gamma)+\hat{\sigma}_\epsilon^2(n-c))^2} \\ \frac{(n-c)\hat{\sigma}_\epsilon^2\text{tr}(\mathbf{MK}_\gamma)}{(\hat{\sigma}_\alpha^2\text{tr}(\mathbf{MK}_\alpha)+\hat{\sigma}_\gamma^2\text{tr}(\mathbf{MK}_\gamma)+\hat{\sigma}_\epsilon^2(n-c))^2} \\ -\frac{(n-c)\hat{\sigma}_\alpha^2\text{tr}(\mathbf{MK}_\alpha)+(n-c)\hat{\sigma}_\gamma^2\text{tr}(\mathbf{MK}_\gamma)}{(\hat{\sigma}_\alpha^2\text{tr}(\mathbf{MK}_\alpha)+\hat{\sigma}_\gamma^2\text{tr}(\mathbf{MK}_\gamma)+\hat{\sigma}_\epsilon^2(n-c))^2} \end{bmatrix},$$

$$\nabla_{prop} = \begin{bmatrix} \frac{\hat{\sigma}_\gamma^2\text{tr}(\mathbf{MK}_\alpha)\text{tr}(\mathbf{MK}_\gamma)}{(\hat{\sigma}_\alpha^2\text{tr}(\mathbf{MK}_\alpha)+\hat{\sigma}_\gamma^2\text{tr}(\mathbf{MK}_\gamma)} \\ -\frac{\hat{\sigma}_\alpha^2\text{tr}(\mathbf{MK}_\alpha)\text{tr}(\mathbf{MK}_\gamma)}{(\hat{\sigma}_\alpha^2\text{tr}(\mathbf{MK}_\alpha)+\hat{\sigma}_\gamma^2\text{tr}(\mathbf{MK}_\gamma)} \\ 0 \end{bmatrix}.$$

Alternatively, we can apply MM algorithm described in [3] to obtain REML estimate of $\boldsymbol{\theta}$ by further assuming the normality $\mathbf{t} \sim \mathcal{N}(\mathbf{t}|\mathbf{Wv}, \sigma_\alpha^2\mathbf{K}_\alpha + \sigma_\gamma^2\mathbf{K}_\gamma + \sigma_\epsilon^2\mathbf{I}_n)$.

##### 2.3.2 IGREX-s

Next, we derive general form of IGREX-s for incorporating covariates for the LD reference $\tilde{\mathbf{X}}$. To obtain $\widehat{\mathrm{PVE}}_{\mathrm{GREX}}$ given by Equation (10) in the main text, we first reorganize (S6) by eliminating the $\sigma_\epsilon^2$ and dividing both sides by $(n-c)^2$:

$$\begin{bmatrix} \frac{\mathrm{tr}((\mathbf{MK}_\alpha)^2)-\frac{\mathrm{tr}^2(\mathbf{MK}_\alpha)}{(n-c)}}{(n-c)^2} & \frac{\mathrm{tr}(\mathbf{MK}_\alpha\mathbf{MK}_\gamma)-\frac{\mathrm{tr}(\mathbf{MK}_\alpha)\mathrm{tr}(\mathbf{MK}_\gamma)}{(n-c)}}{(n-c)^2} \\ \frac{\mathrm{tr}(\mathbf{MK}_\alpha\mathbf{MK}_\gamma)-\frac{\mathrm{tr}(\mathbf{MK}_\alpha)\mathrm{tr}(\mathbf{MK}_\gamma)}{(n-c)}}{(n-c)^2} & \frac{\mathrm{tr}((\mathbf{MK}_\gamma)^2)-\frac{\mathrm{tr}^2(\mathbf{MK}_\gamma)}{(n-c)}}{(n-c)^2} \end{bmatrix} \begin{bmatrix} \sigma_\alpha^2 \\ \sigma_\gamma^2 \end{bmatrix} = \begin{bmatrix} \frac{\mathbf{t}^T\mathbf{MK}_\alpha\mathbf{Mt}}{(n-c)^2} - \frac{\mathrm{tr}(\mathbf{MK}_\alpha)}{(n-c)^3}\mathbf{t}^T\mathbf{Mt} \\ \frac{\mathbf{t}^T\mathbf{MK}_\gamma\mathbf{Mt}}{(n-c)^2} - \frac{\mathrm{tr}(\mathbf{MK}_\gamma)}{(n-c)^3}\mathbf{t}^T\mathbf{Mt} \end{bmatrix}.$$

The terms on the left hand side does not involve $\mathbf{t}$ and thus can be approximated using an LD reference $\tilde{\mathbf{X}}$ [1]. For example, $\frac{\mathrm{tr}(\mathbf{MK}_\alpha^2)-\frac{\mathrm{tr}^2(\mathbf{MK}_\alpha)}{(n-c)}}{(n-c)^2}$ can be well approximated by $\frac{\mathrm{tr}(\tilde{\mathbf{M}}\tilde{\mathbf{K}}_\alpha^2)-\frac{\mathrm{tr}^2(\tilde{\mathbf{M}}\tilde{\mathbf{K}}_\alpha)}{(m-c)}}{(m-c)^2}$, where $\tilde{\mathbf{K}}_\alpha = \sum_{g=1}^G \tilde{\mathbf{X}}_g(\boldsymbol{\mu}_g\boldsymbol{\mu}_g^T + \boldsymbol{\Sigma}_g)\tilde{\mathbf{X}}_g^T$ and $\tilde{\mathbf{M}}$ is the projection matrix corresponding to the covariates associated with the LD reference matrix $\tilde{\mathbf{X}}$. Other terms on the left hand side can be approximated in the same way. Usually, the confounding covariates have been adjusted when calculating the $z$-scores. Thus, the terms in the right hand side can be approximated as follows:

$$\begin{aligned}
\frac{\mathbf{t}^T\mathbf{MK}_\alpha\mathbf{Mt}}{(n-c)^2} &\approx \frac{\mathbf{t}^T\mathbf{K}_\alpha\mathbf{t}}{n^2} \approx \frac{1}{n}\hat{\sigma}_t^2\sum_g \mathbf{z}_g^T(\boldsymbol{\mu}_g\boldsymbol{\mu}_g^T + \boldsymbol{\Sigma}_g)\mathbf{z}_g, \\
\frac{\mathrm{tr}(\mathbf{MK}_\alpha)}{(n-c)^3}\mathbf{t}^T\mathbf{Mt} &\approx \frac{\mathrm{tr}(\mathbf{K}_\alpha)}{n^3}\mathbf{t}^T\mathbf{t} \approx \frac{1}{n}\hat{\sigma}_t^2\mathrm{tr}(\sum_g(\boldsymbol{\mu}_g\boldsymbol{\mu}_g^T + \boldsymbol{\Sigma}_g)\hat{\mathbf{R}}_g), \\
\frac{\mathbf{t}^T\mathbf{MK}_\gamma\mathbf{Mt}}{(n-c)^2} &\approx \frac{\mathbf{t}^T\mathbf{K}_\gamma\mathbf{t}}{n^2} \approx \frac{1}{n}\hat{\sigma}_t^2\sum_{j=1}^M z_j^2, \\
\frac{\mathrm{tr}(\mathbf{MK}_\gamma)}{(n-c)^3}\mathbf{t}^T\mathbf{Mt} &\approx \frac{\mathrm{tr}(\mathbf{K}_\gamma)}{n^3}\mathbf{t}^T\mathbf{t} \approx \frac{1}{n}\hat{\sigma}_t^2.
\end{aligned}$$

Based on these approximations, the estimating equation becomes

$$\begin{bmatrix} \frac{\mathrm{tr}((\tilde{\mathbf{M}}\tilde{\mathbf{K}}_\alpha)^2)-\frac{\mathrm{tr}^2(\tilde{\mathbf{M}}\tilde{\mathbf{K}}_\alpha)}{(m-c)}}{(m-c)^2} & \frac{\mathrm{tr}(\tilde{\mathbf{M}}\tilde{\mathbf{K}}_\alpha\tilde{\mathbf{M}}\tilde{\mathbf{K}}_\gamma)-\frac{\mathrm{tr}(\tilde{\mathbf{M}}\tilde{\mathbf{K}}_\alpha)\mathrm{tr}(\tilde{\mathbf{M}}\tilde{\mathbf{K}}_\gamma)}{(m-c)}}{(m-c)^2} \\ \frac{\mathrm{tr}(\tilde{\mathbf{M}}\tilde{\mathbf{K}}_\alpha\tilde{\mathbf{M}}\tilde{\mathbf{K}}_\gamma)-\frac{\mathrm{tr}(\tilde{\mathbf{M}}\tilde{\mathbf{K}}_\alpha)\mathrm{tr}(\tilde{\mathbf{M}}\tilde{\mathbf{K}}_\gamma)}{(m-c)}}{(m-c)^2} & \frac{\mathrm{tr}((\tilde{\mathbf{M}}\tilde{\mathbf{K}}_\gamma)^2)-\frac{\mathrm{tr}^2(\tilde{\mathbf{M}}\tilde{\mathbf{K}}_\gamma)}{(m-c)}}{(m-c)^2} \end{bmatrix} \begin{bmatrix} \frac{\hat{\sigma}_\alpha^2}{\hat{\sigma}_t^2} \\ \frac{\hat{\sigma}_\gamma^2}{\hat{\sigma}_t^2} \end{bmatrix} = \begin{bmatrix} \frac{\sum_g \mathbf{z}_g^T(\boldsymbol{\mu}_g\boldsymbol{\mu}_g^T+\boldsymbol{\Sigma}_g)\mathbf{z}_g - \mathrm{tr}(\sum_g(\boldsymbol{\mu}_g\boldsymbol{\mu}_g^T+\boldsymbol{\Sigma}_g)\hat{\mathbf{R}}_g)}{n} \\ \sum_{j=1}^M \frac{z_j^2-1}{n} \end{bmatrix}. \tag{S7}$$

Solving this equation and plugging the obtained $\frac{\hat{\sigma}_\alpha^2}{\hat{\sigma}_t^2}$ into Equation (10) give the estimate of $\mathrm{PVE}_{\mathrm{GREX}}$. When the summary data is from meta-analysis of multiple GWASs, the sample size for each SNP may be different. To handle this heterogeneity in sample size, the right hand side of (S7) is further substituted by

$$\left[\sum_g \tilde{\mathbf{z}}_g^T(\boldsymbol{\mu}_g\boldsymbol{\mu}_g^T + \boldsymbol{\Sigma}_g)\tilde{\mathbf{z}}_g - \sum_g\left[\mathrm{tr}((\boldsymbol{\mu}_g\boldsymbol{\mu}_g^T + \boldsymbol{\Sigma}_g)\hat{\mathbf{R}}_g)(\tfrac{1}{M_g}\textstyle\sum_j^{M_g}\tfrac{1}{n_j})\right],\ \ \textstyle\sum_{j=1}^M \tfrac{z_j^2-1}{n_j}\right]^T,$$

where $\tilde{\mathbf{z}}_g = [\mathbf{z}_{g,1}/\sqrt{n_1}, ..., \mathbf{z}_{g,j}/\sqrt{n_j}, ..., \mathbf{z}_{g,M_g}/\sqrt{n_{M_g}}]^T$. The standard errors of estimated PVE are calculated by block-wise jackknife where the $z$-scores are divided into $1,704$ approximately independent blocks according to the LD scores of corresponding SNPs [4]. To guarantee the independence between blocks, only the genes whose local SNPs all lie inside an LD block are included for calculation. When there are large overlaps between gene coverage and LD block cut-off, may genes can be excluded from the calculation. In this case, the jackknife approach tends to slightly over-estimate the standard error (See Supplementary Fig. 7).

#### 2.4 Application to metabolite traits

Metabolic phenotypes serve as important intermediate traits in high level biological processes. To understand the role of gene regulation in the genetics of such traits, we applied IGREX-s to a summary level data set of circulating metabolites [5], which was comprised of meta-analysis of 123 metabolites. We focused our analysis on the 21 metabolites that were highly heritable (estimated $h_t^2 > 10\%$) including glycine, various features of HDL, LDL, very low-density lipoprotein (VLDL) and intermediate-density lipoprotein (IDL) and other polyunsaturated fatty acids (otPUFA). The distributions of $\mathrm{PVE_{GREX}}/h_t^2$ estimates in different tissues are given in Supplementary Fig. 16a. The median values of percentage estimates are higher than 10% in 6 out of the 48 tissues and only higher than 15% in liver and spinal cord (cervical c-1). According to the estimated values shown in the heat map of Supplementary Fig. 16b, we can see that the features associated with IDL, LDL and VLDL have estimated $\mathrm{PVE_{GREX}}/h_t^2$ around 20% in liver and 16% in spinal cord, suggesting that they are more related to the GREX effects in these two tissues. On the other hand, there is no signal of GREX components detected under the nominal level 0.05 in any GTEx tissue for HDL associated features or glycine. We note that the estimated values for LDL are not significantly different from the ones observed in NFBC analysis, implying the consistency between the two studies.

#### 2.5 Details of trans-eQTLs and alternative splicing analysis

The trans-eQTLs used in analysis are reported by an independent study, the eQTLGen consortium. After matching with the SNPs in both the GWAS (or the LD reference panel) and the eQTL reference panel, around 20,000 trans comprised of 4,000 genes and 1600 SNPs were incorporated in the model for protein traits and the two schizophrenia datasets. In analyzing HDL, 8,072 trans-associations from 2,817 genes and 663 SNPs were included. In diseases

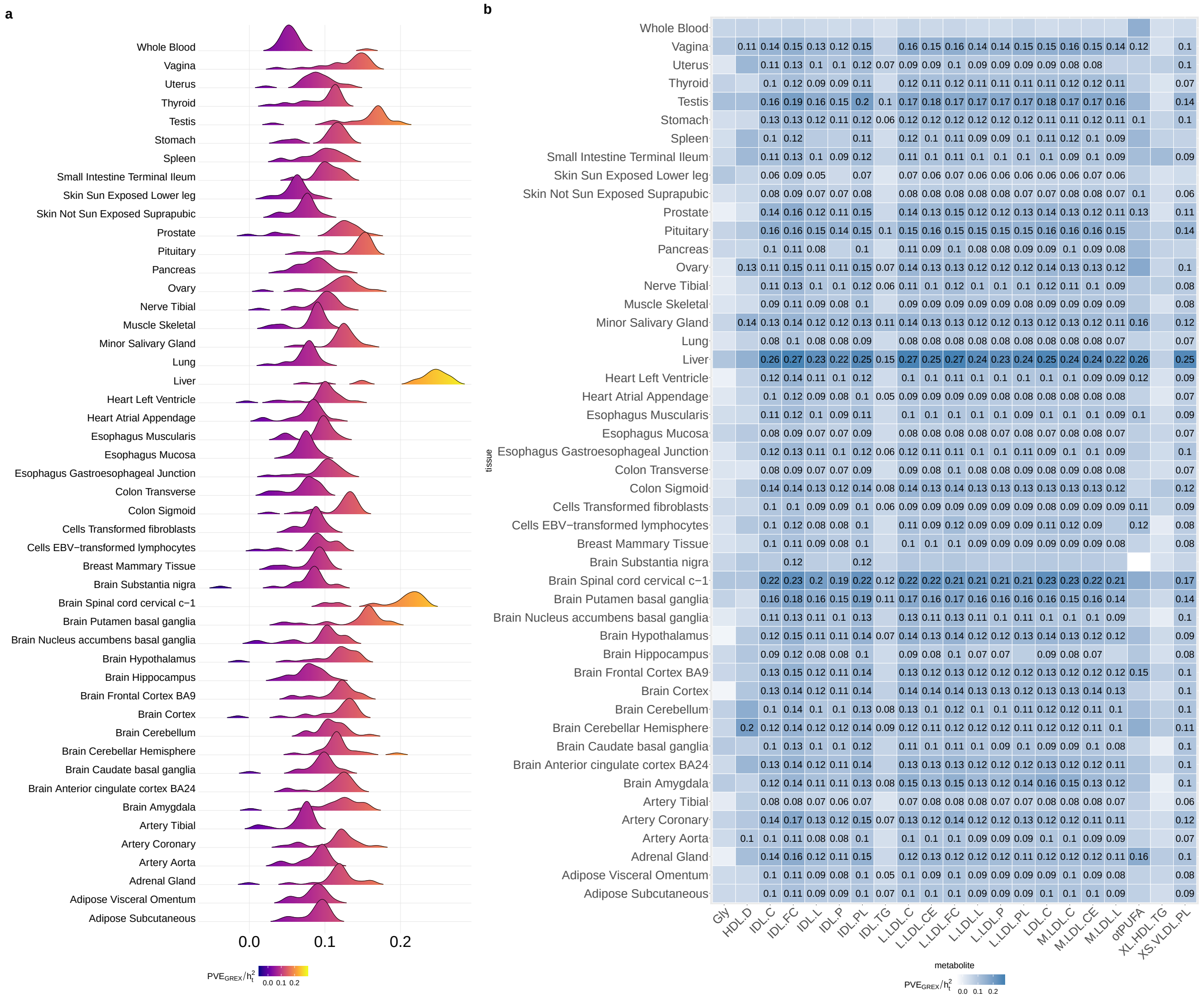

**Supplementary Figure 16:** $\widehat{\text{PVE}}_{\text{GREX}}/h_t^2$ for 21 circulating metabolites. (**a**) The distributions of estimated $\text{PVE}_{\text{GREX}}/h_t^2$ in different tissues. (**b**) The heat map of estimated $\text{PVE}_{\text{GREX}}/h_t^2$. Entries that are significant at nominal level (0.05) are labeled with their estimated values.

from WTCCC dataset, around 5,700 trans-associations from 2,300 genes and 350 SNPs were considered.

In the analysis of sQTL data, we used the alternative splicing data from GTEx as reference and applied IGREX-MoM to four representative trait-tissue pairs: LDL-liver, TC-liver, SCZ-amygdala and SCZ-cerebellar hemisphere. The LDL and TC datasets were from the NFBC study. SCZ2, the dataset with the largest sample size, was used for SCZ. After matching with the GWAS, 58,272 splicing events were analyzed for LDL and TC while 57,509 splicing events were analyzed for SCZ.
